# Convergence, stability, and thermal adaptation in the rubisco enzyme in plants

**DOI:** 10.1101/2025.10.08.681247

**Authors:** Arthur Leung, Belinda S. W. Chang, Rowan F. Sage

**Affiliations:** Department of Ecology and Evolutionary Biology, University of Toronto, 25 Willcocks Street, Toronto, ON M5S 3B2, Canada; Department of Cell and Systems Biology, University of Toronto, 25 Harbord Street, Toronto, ON M5S 3G5, Canada

**Keywords:** biochemical adaptation, growing season temperature, homology modelling, phylogenetic comparative methods, protein stability, rubisco

## Abstract

Enzymes are thought to be tuned to perform similarly in different thermal regimes. Whether the photosynthetic enzyme ribulose-1,5-bisphosphate carboxylase/oxygenase (rubisco) follows similar rules, especially when considering evolutionary history, is uncertain. The molecular, structural, and ecological factors of the rubisco large subunit (RbcL) were examined in four plant clades: wood ferns, pines, sea lavenders, and viburnums. Using *rbcL* gene sequences, codon evolutionary models were used to test for positive and divergent selection and convergent evolution. Protein structure modeling was performed to predict side chain changes and protein stability. Phylogenetic comparative methods were used to examine the relationship between protein stability to growing season temperature. All four clades showed significant evidence of positive selection, with multiple convergent substitutions predicted to alter side chain polarity and interactions with the solvent. In viburnums, biome transition rates were dependent on amino acid substitution, with positive selection was concentrated in cold temperate and cloud forest clades. Rubiscos with higher stability occurred in species from warmer environments. However, this correlation was weaker after correcting for phylogeny. These analyses support a hypothesis that RbcL evolution is influenced by both environmental tuning and evolutionary history.

## Introduction

The diverse range of environmental conditions on the planet are thought to contribute to the evolution of enzyme performance, leading to adaptations that tune function such as catalytic rate and substrate affinity. These properties are represented by the turnover rate per active site (*k*_cat_) and the inverse of the Michaelis constant (*K*_m_). Enzyme stability and flexibility are key determinants of turnover rate and affinity. Flexible enzymes bind substrate and release product faster but are more prone to denaturing or adopting low-affinity conformations (Fields, 2001; Fields and Houseman, 2004). Variations in temperature disrupt this stability-flexibility balance. At supraoptimal temperatures, non-covalent interactions (e.g., hydrogen bonds) that stabilize the enzyme are weakened, initially leading to elevated reaction rates but decreased affinity; in contrast, at suboptimal temperatures, non-covalent interactions are strengthened, reducing time spent in low-affinity conformations, but inhibiting catalytic motions that contribute to greater *k*_cat_ (Fields, 2001).

The temperature response of reaction rate is typically exponential and doubles with every 10 °C rise in temperature (Scholander et al., 1953). However, in a comparative context, proteins in different species do not tend to follow this relationship. For example, a fish that typically lives at 0 °C does not have a metabolic rate that is 20 times slower than one that typically lives at 40 °C (Fields et al., 2015). Evolutionary selection drives distinct adaptations in organisms from different thermal habitats such that stability and flexibility is optimized for their environment (Hochachka and Somero, 2002). As shown for lactate dehydrogenase (LDH) and malate hydrogenase (MDH), cold-adapted enzymes are more flexible and less stable than warm-adapted orthologs, enabling similar performance (Fields and Somero, 1998; Dong and Somero, 2009; Liao et al., 2017; Dong et al., 2018). Thus, protein evolution allows organisms to escape the “tyranny” of the exponential temperature response (Bullock, 1955).

While LDH and MDH are classic examples of thermal adaptation in enzymes, ribulose-1,5-bisphosphate (RuBP) carboxylase/oxygenase (rubisco), which catalyzes CO_2_ fixation in photosynthesis, presents additional challenges due to its dual-substrate specificity. The carboxylation of one RuBP molecule produces two phosphoglycerate molecules, which is used in the photosynthetic carbon reduction cycle to produce glucose. Oxygen competitively inhibits RuBP carboxylation, and each RuBP oxygenation event produces one phosphoglycerate and one phosphoglycolate molecule (Bowes and Ogren, 1972). Phosphoglycolate is toxic and must be recycled into phosphoglycerate, which consumes energy and releases previously fixed CO_2_ in the process. Importantly, the *k*_cat_ and *K*_m_ for CO_2_ and O_2_ (*k*_cat_^c^, *K*_c_ and *k*_cat_°, *K*_o_, respectively) do not scale uniformly to temperature (Jordan and Ogren, 1984). However, variation in *k*_cat_^c^ and CO_2_ specificity factor (*S*_c/o_) across species indicates evolutionary flexibility (Jordan and Ogren, 1981; Galmés et al., 2016). In an extreme case, CO_2_ specificity varies by an order of magnitude between *Rhodospirillum rubrum* and land plants (Jordan and Ogren, 1981). Rubisco small subunit (RbcS) isoforms can further modulate kinetics in plants, adding an another layer of complexity (Lin et al., 2020; Cavanagh et al., 2022). Whether these patterns are consistent with broader trends in enzyme evolution, such as those in LDH and MDH (Hochachka and Somero, 2002), remains unresolved.

The complexity of rubisco kinetics becomes more apparent when considering how variation in CO_2_ availability, shaped by environmental pressures and photosynthetic pathways, influences its evolution. Plants with CO₂-concentrating mechanisms (CCMs), such as C₄ and CAM species, operate at higher plastidial CO₂ concentrations than C₃ plants, which has driven rubisco adaptation toward higher *k*_cat_^c^ but lower *K*_c_ (Yeoh et al., 1980, 1981; Iñiguez et al., 2020). High CO_2_ affinity is less important than enzyme velocity when the CO_2_ availability at the active sites is high, and thus these kinetic differences have convergently evolved multiple times (Christin et al., 2008). In plants that lack CCMs, rubisco adaptation is constrained by the relative rates of carboxylation to oxygenation and the trade-off between *k*_cat_^c^ and *S*_c/o_ (Lorimer and Andrews, 1973; Tcherkez et al., 2006; Savir et al., 2010). It is thought that better differentiation of CO_2_ and O_2_ transition states strengthens the binding with transition intermediates, slowing subsequent steps in the reaction (Tcherkez et al., 2006). For instance, C_3_ plants in warm, saline, or arid environments, where stomatal closure limits diffusion of CO_2_ substrate to the active site, tend to have rubiscos with lower *k*_cat_^c^ but higher *S*_c/o_ or lower *K*_c_ (Sage, 2002; Galmés et al., 2005). However, this *k*_cat_-*K*_c_ trade-off is weaker in recent, larger datasets (Flamholz et al., 2019), which would raise questions regarding if and how evolutionary history can shape RbcL evolution.

A comparative phylogenetic approach offers a powerful means of understanding the evolution of rubisco structure and function in relation to temperature. It can be used to identify sites in the protein that were the targets of positive selection, and to locate amino acid shifts onto specific lineages of the phylogeny in distant relatives (Kapralov and Filatov, 2007; Galmés et al., 2014a; Hermida-Carrera et al., 2016; Orr et al., 2016; Sharwood et al., 2016) and within genera (Kubien et al., 2008; Galmés et al., 2014a; Capó-Bauçà et al., 2023). Although amino acid changes in rubisco can be linked to shifts in enzymatic function, the connection between amino acid variation and its broader ecological context is less frequently explored (but see Hermida-Carrera et al., 2017). Phylogenetic comparative analyses are thus useful, because evolutionary relatedness between proteins can confound the interpretation of rubisco evolution (Tcherkez et al., 2006; Savir et al., 2010; Bouvier et al., 2021, 2024; Tcherkez and Farquhar, 2021; Bouvier and Kelly, 2023). For instance, the relationship of *k* ^c^ with *K*_c_ may differ when compared across broad versus narrow taxonomic scales (Galmés et al., 2014b). Thus, evolutionary history must be explicitly accounted for when correlating amino acid sequence with different thermal niches (Felsenstein, 1985; Pagel, 1994).

This framework allows us to account for phylogenetic non-independence and to better resolve whether observed trait correlations are adaptive or simply reflect the degree of relatedness. For instance, distantly relatedly species may differ in the genetic background other than RbcL, which could confound differences in *k*_cat_^c^ and *K*_c_ (Hermida-Carrera et al., 2016) On the other hand, examining closely related species without correcting for relatedness can lead to misleading conclusions about adaptation, since similarities may be due to shared ancestry rather than adaptation to environmental pressures (Bouvier et al., 2021). Based on this rationale, we selected phylogenetically disparate plant genera each spanning multiple habitats: wood ferns (*Dryopteris*), sea lavenders (*Limonium*), pines (*Pinus*), and viburnums (*Viburnum*). We asked the following questions: (1) Is adaptive evolution responsible for RbcL diversity? (2) Is RbcL diversity associated with the thermal environment of vascular plant species? (3) How do these patterns hold up when correcting for phylogenetic history?

By using phylogenetics to examine structural and evolutionary patterns in tandem, we aim to evaluate how molecular changes in rubisco explain its adaptation. We hypothesized that convergent amino acid substitutions would be consistent with an increased the number of non-covalent interactions in the holoenzyme that enhance stability in warm or dry climates. By integrating analyses of selection, convergence, stability, and correlated evolution, our study provides insight into the evolutionary and ecological basis of rubisco diversity.

## Materials and methods

### Taxon sampling and analytical approach

We used a model clade approach (Donoghue and Edwards, 2019; Mabry et al., 2024). Patterns of positive selection on a large phylogeny consisting of plants and algae (Kapralov and Filatov, 2007) may differ from those in closely-related species (Galmés et al., 2014a). Sparse taxon sampling of distantly related species can lead to the misinterpretation of other evolutionary change, such as that in rubisco small subunit (RbcS) and rubisco activase (Rca), with changes associated with the focal trait (Heyduk et al., 2019). We chose four phylogenetically distant genera that each were speciose and had well-resolved phylogenies, such that each can be examined as separate examples of evolution, and such that effects of saturation are minimized.

The four genera, *Dryopteris*, *Limonium*, *Pinus*, and *Viburnum,* were chosen. Tests of saturation showed minimal saturation in *Dryopteris*, *Pinus*, and *Viburnum* (Table S 1). *Limonium* showed evidence of saturation, which could lead to overestimation of positive selection; thus, the significance of the results may be weakened.

### Gene sequences and alignments

*rbcL* gene sequences from 621 vascular plant accessions were downloaded from GenBank: 185 *Dryopteris*, 184 *Limonium*, 104 *Pinus*, and 148 *Viburnum* accessions (see Data Availability for accession numbers). Separately for each genus, sequences were aligned using MUSCLE v5 (Edgar, 2022) and then manually inspected in MEGA v11 (Tamura et al., 2021). To facilitate homology modelling, the sequences were aligned with a spinach *rbcL* sequence (GenBank: NC 002202), which was later removed from the alignment. Species-level phylogenetic trees for *Pinus* and *Viburnum* were downloaded from supplementary data of respective papers (Jin et al., 2021; Landis et al., 2021). The *Dryopteris* and *Limonium* phylogenies were provided by the lead authors (Sessa et al., 2017; Koutroumpa et al., 2021); both lead authors granted written permission to make these files available in the data repository (see Data Availability).

### Codon evolutionary analyses

Random sites models were used to analyze codon evolution at the site level using CodeML in ‘PAML’ v4.10 (Yang, 2007). Codon model (M) 0, M1a, M2a, M3, M7, and M8 were used with codon frequencies empirically determined from the *rbcL* sequences. For each genus, the best fitting codon model with the lowest Akaike Information Criterion (AIC) value was used to reconstruct of the ancestral state at the root of the phylogeny. Likelihood-ratio test results of M1 vs. M0, M2a vs. M1a, M3 vs. M0, and M8 vs. M7 are reported in Table 1. Bayes empirical Bayes (BEB) analysis determined the probability that each codon site was under positive selection; codon sites with a BEB posterior probability (*P*) > 0.95 were considered under positive selection.

**Table 1.** Likelihood ratio tests of PAML codon models in each genus (Yang, 2007). For models M0–M3, *ω*_0_–*ω*_2_ values are shown representing *d*_N_/*d*_S_ values for each site class, along with proportion of sites for each site class. For models M7 and M8, α and β describe the shape of the beta distribution, and *ω*_P_ describes the *d*_N_/*d*_S_ value for the positively selected site class. n.p., number of parameters; AIC, Akaike Information Criterion; ln*L*, log-likelihood; κ, transition/transversion ratio; *χ*^2^, chi-squared; d.f., degrees of freedom; *p*, *p*-value of likelihood ratio test.

| Genus | Model | n.p. | AIC | $\ln L$ | $\kappa$ | $\omega_0/\alpha$ | $\omega_1/\beta$ | $\omega_2/\omega_P$ | $\chi^2$ | d.f. | $p$ |
| --- | --- | --- | --- | --- | --- | --- | --- | --- | --- | --- | --- |
| *Dryopteris* | M0 | 369 | 17871 | -8566.5 | 6.92 | 0.1 |  |  |  |  |  |
| · | M1a | 370 | 17163 | -8211.4 | 6.69 | 0 (84.5%) | 1 (15.5%) |  | 355.1 | 1 | <0.001 |
| · | M2a | 372 | 16867 | -8061.6 | 7.45 | 0 (88.1%) | 1 (7.2%) | 42.0 (4.7%) | 149.8 | 2 | <0.001 |
| · | M3 | 373 | 16857 | -8055.6 | 7.8 | 0 (4.8%) | 2.0 (90.3%) | 46.5 (4.9%) | 510.9 | 4 | <0.001 |
| · | M7 | 370 | 16863 | -8209.8 | 6.73 | 0.078 | 0.369 |  |  |  |  |
| · | M8 | 372 | 16922 | -8088.8 | 7.55 | 0.067 | 0.427 | 41.5 (0.045%) | 121 | 2 | <0.001 |
| *Limonium* | M0 | 367 | 7282 | -3274.2 | 2.11 | 0.1 |  |  |  |  |  |
| · | M1a | 368 | 7004 | -3134.0 | 1.92 | 0 (93.5%) | 1 (6.5%) |  | 140.2 | 1 | <0.001 |
| · | M2a | 370 | 6926 | -3093.1 | 2.1 | 0 (93.4%) | 1 (5.2%) | 7.1 (1.4%) | 40.9 | 2 | <0.001 |
| · | M3 | 371 | 6964 | -3110.8 | 2.07 | 0 (97.7%) | 5.5 (2.2%) | 999 (0.1%) | 163.4 | 4 | <0.001 |
| · | M7 | 368 | 6922 | -3142.8 | 1.92 | 0.005 | 0.048 |  |  |  |  |
| · | M8 | 370 | 7019 | -3139.7 | 1.94 | 0.005 | 0.048 | 999 (0.001%) | 3.1 | 2 | 0.21 |
| *Pinus* | M0 | 207 | 8040 | -3812.8 | 2.5 | 0.4 |  |  |  |  |  |
| · | M1a | 208 | 7661 | -3622.5 | 2 | 0 (91.1%) | 1 (8.9%) |  | 380.6 | 1 | <0.001 |
| · | M2a | 210 | 7667 | -3623.5 | 2 | 0 (90.9%) | 1 (8.9%) | 999 (0.1%) | -1.8 | 2 | 1.00 |
| · | M3 | 211 | 7453 | -3515.6 | 2.38 | 0 (94.6%) | 4.2 (4.0%) | 17.8 (1.4%) | 594.5 | 4 | <0.001 |
| · | M7 | 208 | 7663 | -3622.8 | 2.01 | 0.005 | 0.047 |  |  |  |  |
| · | M8 | 210 | 7456 | -3517.9 | 2.38 | 0.005 | 0.051 | 11.3 (0.030%) | 209.6 | 2 | <0.001 |
| *Viburnum* | M0 | 295 | 8039 | -3724.6 | 1.14 | 1.0 |  |  |  |  |  |
| · | M1a | 296 | 7622 | -3515.0 | 0.8 | 0 (90.1%) | 1 (9.9%) |  | 419.2 | 1 | <0.001 |
| · | M2a | 298 | 7240 | -3322.1 | 1.06 | 0 (87.9%) | 1 (7.4%) | 18.9 (4.6%) | 385.7 | 2 | <0.001 |
| · | M3 | 299 | 7220 | -3311.0 | 1.12 | 0 (93.1%) | 8.1 (4.5%) | 32.5 (2.5%) | 827.1 | 4 | <0.001 |
| · | M7 | 296 | 7236 | -3603.9 | 0.97 | 0.005 | 0.005 |  |  |  |  |
| · | M8 | 298 | 7241 | -3322.5 | 1.07 | 0.005 | 0.049 | 19.7 (0.046%) | 563 | 2 | <0.001 |

### Protein convergence analyses

Using ‘CSUBST’ v1.4.0 (Fukushima and Pollock, 2023), ancestral gene sequences were reconstructed to locate branch pairs at which convergent non-synonymous and synonymous substitutions occurred (any amino acid to a specific amino acid) and calculate the rate at which they occur for each codon site (hereafter, *d*_Nc_ and *d*_Sc_, respectively). ‘CSUBST’ internally uses ‘IQTREÈ to determine the best codon substitution model for ancestral state reconstruction (Minh et al., 2020; Fukushima and Pollock, 2023). In contrast to commonly used methods to examine *rbcL* convergence, this method corrects for phylogenetic errors due to introgression (Fukushima and Pollock, 2023). Introgression has been shown to affect *rbcL* evolution (Yao et al., 2019) and would similarly affect both the *d*_Nc_ and *d*_Sc_, so not accounting for the *d*_Sc_ would overestimate the rate of convergent codon evolution. In contrast, convergent amino acid substitutions that arise from positive selection should increase the *d*_Nc_ relative to the *d*_Sc_. Thus, convergent amino acid substitutions were defined as a substitution at the same site in a branch pair any amino acid (empirical Bayesian *P* > 0.95) to the same amino acid, with a *d*_Nc_/*d*_Sc_ > 1 (Fukushima and Pollock, 2023).

### Amino acid- and codon-level associations with biome transitions in Viburnum

For *Viburnum*, discrete biome scorings were obtained by binarizing published data (Landis et al., 2021). Discrete representations of thermal niche based on expert observations were unavailable for the other three genera at the time of this work. *Viburnum* species occur in warm temperate forests, cold temperate forests, cloud forests, and tropical forests. The genus first diversified in warm temperate forests and later diversified into the other biomes (Landis et al., 2021).

To further examine the associations of codon evolution with transitions between biomes, clade models C and D (CmC, CmD) in CodeML were used. These models test for divergence in *d*_N_/*d*_S_ between clades ‘PAML’ v4.10 (Yang, 2007). Clade models allow for one class of codon sites to vary in *d*_N_/*d*_S_ (*ω*) between preassigned clades (e.g., clades occupying different biomes).

Compared to branch-site models, which test for rapid positive selection on specific branches, clade models can test for long-term selection across clades (Chang et al., 2012). CmC allows three classes of sites with 0 < *ω*_0_ < 1, *ω*_1_ = 1, and *ω*_2_ > 0, with *ω*_2_ varying between clades. CmD relaxes the assumptions of the first two classes, allowing *ω*_0_, *ω*_1_ > 0 (Bielawski and Yang, 2004). The null models of CmC and CmD are M2a_rel (Weadick and Chang, 2012) and M3, both of which constrain *ω*_2_ to be identical across clades. Near-identical parameter estimations resulted from different initial values of transversion/transition ratio (0.3, 10) and/or *ω* (1.3, 10) (see Data Availability).

The dependence of biome evolution on amino acid evolution across the *Viburnum* phylogeny (Landis et al., 2021) was tested by fitting models of evolution in which they were dependent or independent. Binary character evolution models (Pagel, 1994) were fitted using the ‘fitPagel’ function in ‘phytools’ v1.9-9 in R (Revell, 2012). The Pagel (1994) test provides a conservative test of correlated evolution by counting independent transitions rather than inflating sample size with phylogenetically non-independent species observations, making it appropriate for detecting coevolutionary patterns. The models can accommodate cases where biome or amino acid transitions occur before or after the other. Tests evaluated the dependence of biome transition rates on amino acid transition rates and vice versa, alternately designating amino acid or biome as the independent variable; models were selected based on the lowest AIC values. Likelihood ratio tests compared the model in which biome and amino acid evolve independently versus a model in which the two characters evolve dependently.

### Homology modelling

The effects of amino acid substitution on non-covalent interactions were visualized using ‘PyMol’ v3.0.4 (Schrodinger, 2015). Point mutations that add or remove oxygen or sulfur atoms were introduced to the large subunit of a 1.6 Å spinach rubisco structure containing residues 9– 475 in the large subunit (PDB: 8RUC). The lowest energy rotamer was used. Substitutions were assessed based on whether they altered the number of non-covalent interactions within 3.7Å of the amino acid residue.

### Protein stability modelling

The free energy of protein folding (Δ*G*_fold_) of the spinach rubisco structure was predicted at 298 K using ‘FoldX’ v5.1 (Delgado et al., 2019). ‘FoldX’ is widely used for protein stability predictions and its predictions reliably reproduce experimental data (Khan and Vihinen, 2010; Buß et al., 2018), but it is important to interpret modelled results with caution. Following Studer et al. (2014), heteroatoms that are not well parameterized in FoldX (carboxyarabinitol bisphosphate and Mg^2+^) were removed from the structure, with interacting protein residues constrained in their conformation. The residues interacting with these heteroatoms according to Andersson (1996) were constrained in space as done previously (Studer et al., 2014). Using ‘RepairPDB’ in ‘FoldX’, high energy residues were adjusted to its lowest energy rotamer (Studer et al., 2014). Modeling mutations on a single reference structure is standard practice for comparative protein evolution studies (Studer et al., 2014) and is appropriate here because most positively selected sites occur at surface-exposed positions (Figure 1) where local structural context is less constrained than in the catalytic core.

**Figure 1.**
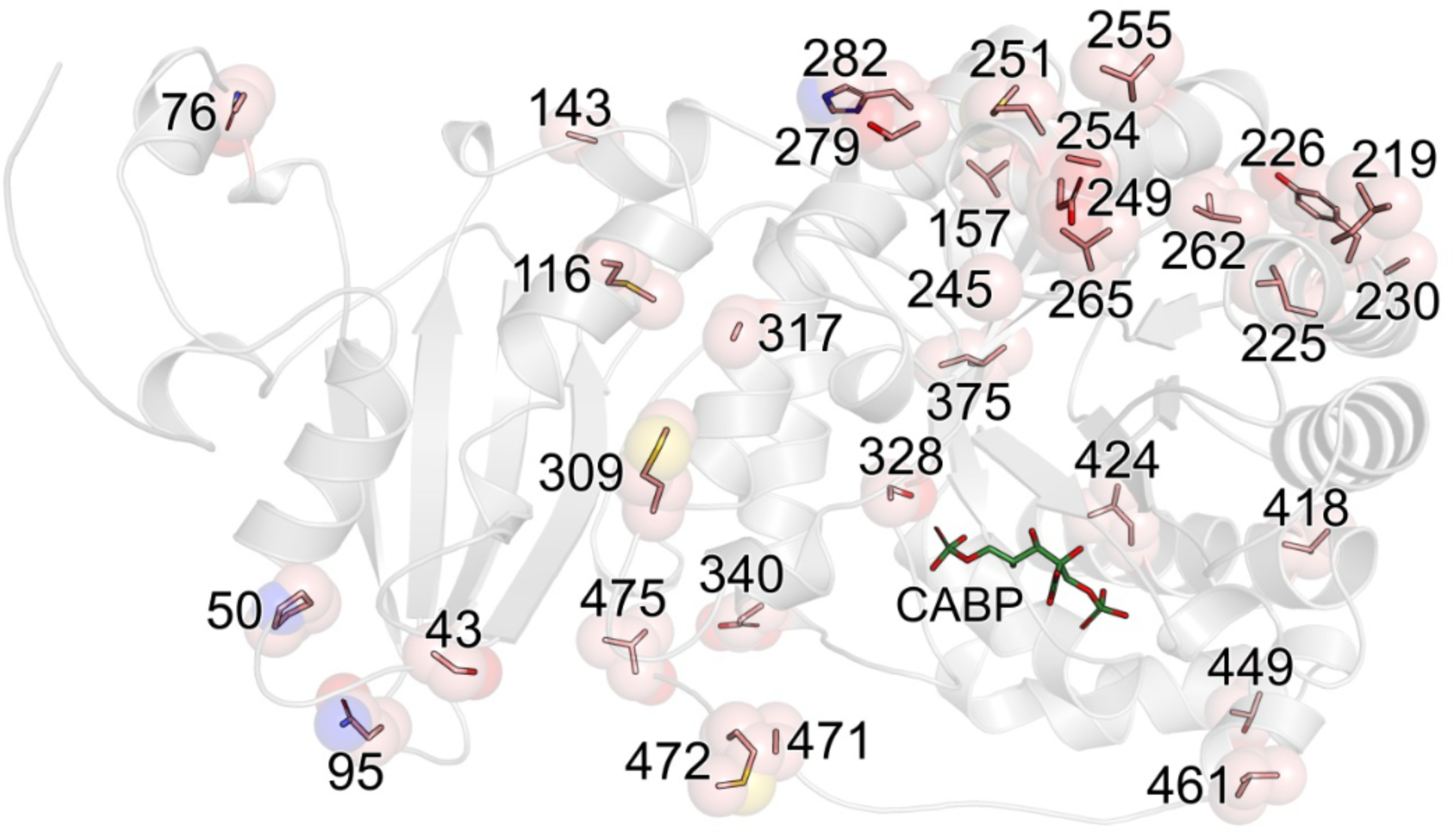
Sites under positive selection and/or convergent evolution in *Dryopteris*, *Limonium*, *Pinus*, and/or *Viburnum* overlayed on a published crystal structure of rubisco large subunit protein bound to transition-state analog carboxyarabinitol bisphosphate (CABP) from spinach (PDB: 8RUC) (Andersson, 1996).

For each substitution at sites under positively selection and/or convergent evolution, all large subunits in the rubisco holoenzyme were mutated, with one substitution at a time, using ten replicate runs of ‘BuildModel’ in ‘FoldX’. The free energies of protein folding in the wildtype (Δ*G*_fold,wt_) and the mutant protein (Δ*G*_fold,mut_) were predicted. The reported ΔΔ*G*_fold_ = Δ*G*_fold,wt_ − Δ*G*_fold,mut_ values were divided by eight to standardize the values to each large subunit (Studer et al., 2014). For each amino acid substitution, the wildtype residue was set to the ancestral residue in the relevant clade, while the mutant residue was set to the derived residue. For sites where the derived residue matched the residue in spinach, the ΔΔ*G* was assumed to be path-independent (e.g., ΔΔ*G*_N76→S_ = −ΔΔ*G*_S76→N_) (Studer et al., 2014). For sites where neither ancestral nor derived residues match the residue in spinach, the ΔΔ*G* values were assumed to be additive and path-independent (e.g., ΔΔ*G*_A328→G_ = ΔΔ*G*_S328→G_ − ΔΔ*G*_S328→A_) (Studer et al., 2014).

A separate analysis predicted the ΔΔ*G* for each species relative to the spinach homology model. For all RbcL gap-free sequences in this study (*n* = 399 species; *n* = 174 unique protein sequences; mean sequence coverage = 92% of the full coding sequence), the spinach crystal structure was mutated to each unique sequence in ‘FoldX’. For the remaining 8% of missing data at the N- and C-termini, the spinach wild-type sequence was used. The analysis was parallelized locally using GNU Parallel (Tange, 2023).

### Growing season temperature estimation

Selection on rubisco performance was assumed to be strong during warm and wet months, when stomata are open and leaves are photosynthetically active. We used interpolated climate data from WorldClim as a standardized, reproducible metric of thermal environment, following established protocols where species-specific microclimate data are unavailable (Fick and Hijmans, 2017; Wright et al., 2017; He et al., 2020). We downloaded geographical coordinates corresponding to species occurrences for each genus (GBIF, 2023, 2024a, b, c). Incomplete or fuzzy species name searches on GBIF were manually inspected. ‘CoordinateCleaner’ was used to exclude duplicate coordinates, (0,0) coordinates, points with identical latitude and longitude values, oceanic points, points overlapping with biodiversity institutions, points near country capitals, and points near country centroids. For species with three or more occurrences, monthly temperature and aridity index values were downloaded from WorldClim and Global Aridity Index and Potential Evapotranspiration Database (Fick and Hijmans, 2017; Zomer et al., 2022). The growing season was defined the months with monthly mean temperature ≥ 5°C and aridity index > 0.05 (Wright et al., 2017; He et al., 2020). The mean temperature of the warmest month in the growing season was used.

### Phylogenetic generalized least squares regressions

To examine the evolution of RbcL stability across genera, a backbone tree was created with crown ages of *Dryopteris*, *Limonium*, *Pinus*, and *Viburnum* with divergence times from Timetree.org; previously published species trees (Sessa et al., 2017; Jin et al., 2021; Koutroumpa et al., 2021; Landis et al., 2021) were then grafted onto the backbone tree (see Data Availability). Statistical tests were performed using phylogenetic generalized least squares regressions under a Brownian motion model implemented in ‘apè with post-hoc tests performed using estimated marginal means implemented in ‘emmeans’ (Searle et al., 1980; Paradis and Schliep, 2019).

## Results

### Pervasive positive selection in rbcL

To test for positive selection at the codon level in the *rbcL* gene, we used PAML to fit random sites models of codon evolution to the alignment and phylogeny corresponding to each genus (Yang, 2007). Null models in which 0 ≤ *d*_N_/*d*_S_ ≤ 1 (M1a and M7) were compared to corresponding models with an additional class of codons under positive selection (*d*_N_/*d*_S_ > 1; M2a and M8, respectively). We also fitted a model that imposes no specific constraints on three categories of *d*_N_/*d*_S_ (M3), as a potential model to use for ancestral state reconstruction.

As summarized in Table 1, the likelihood ratio tests of random sites models show at least one or both of the positive selection models (M2a and M8) were found to be a significantly better fit than the respective null models (*p* < 0.001), providing evidence for positive selection in *rbcL* across all four genera. A Bayes empirical Bayes (BEB) procedure was also used to calculate the posterior probability (*P*) of each site being under positive selection. Sites with *P* > 0.95 were categorized as under positive selection (Table 1; Figure 2); these sites had *d*_N_/*d*_S_ values ranging from 6 to 10, indicating a high rate of amino acid substitution. The number of sites under positive selection were 5 in *Dryopteris*, 5 in *Limonium*, 14 in *Pinus*, and 19 in *Viburnum* (Figure 2). As visualized on the crystal structure, many of these sites were located on the surface of the large subunit and few were around the active site (Figure 1).

**Figure 2.**
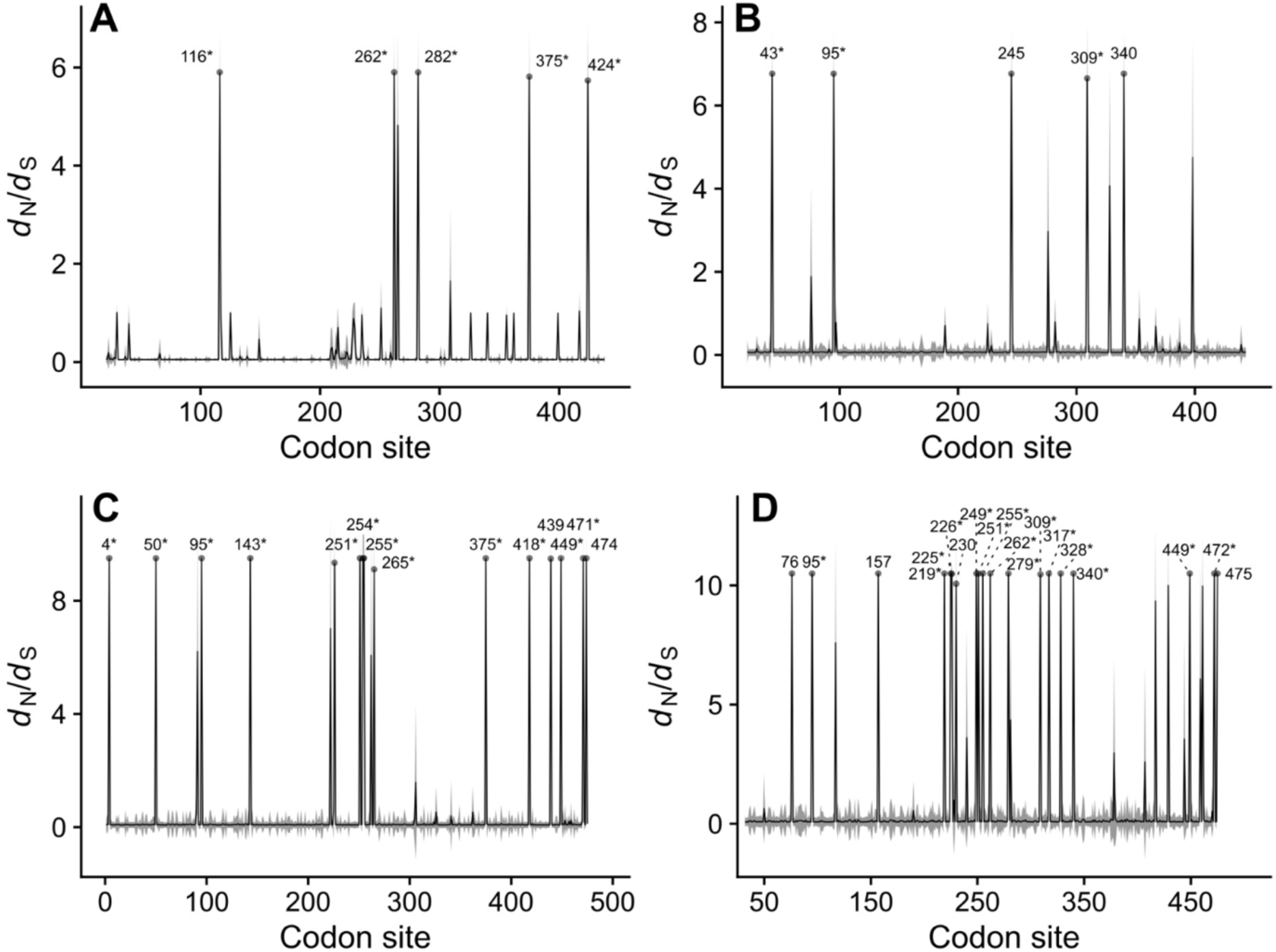
The non-synonymous substitution rate (*d*_N_): synonymous substitution rate (*d*_S_) ratio at each codon site of rubisco large subunit in (A) *Dryopteris*, (A) *Limonium*, (C) *Pinus*, and (D) *Viburnum*. Symbols and labels indicate sites under positive selection (Bayes empirical Bayes *P* > 0.95). Asterisks (*) indicate sites under convergent evolution (empirical Bayes *P* > 0.95).

### High levels of protein convergence within genera

While positive selection suggests adaptive evolution, convergent evolution would indicate that amino acid substitutions occurred due to similar selective pressures rather than shared ancestry. This was tested using ancestral codon reconstruction at each branch and estimating a convergent *d*_N_/*d*_S_ (i.e., *d*_Nc_/*d*_Sc_; see Materials and Methods) across all branch pairs. Using the *d*_Sc_ corrects for introgression and reduces the likelihood of overestimating convergence.

Using this method, most of the sites under positive selection were also identified to be under convergent evolution (empirical Bayes *P* > 0.95). Each convergent substitution typically occurred across two to six branches (Table S 3). However, parallel evolution in rubisco at the codon level was sometimes extensive; for instance, A262→V occurred across 19 different branches in the *Dryopteris* phylogeny (Table S 3). Additionally, at one site, we observed convergence towards two different derived amino acids, e.g., A/C449→S and C/S449→A in *Pinus* (Table S 3). The high probability of protein convergence at multiple sites provides further support for the adaptive evolution of the *rbcL* gene.

### Codon and amino acid substitutions were correlated with biome transitions in Viburnum

Clade models C and D (CmC, CmD) were used to test for divergences in *d*_N_/*d*_S_ between clades in viburnums (Bielawski and Yang, 2004). This genus of plants first diversified in warm temperate (lucidophyllous) forests and subsequently underwent multiple independent radiations into cold temperate and tropical forests, as well as a single radiation into cloud forests (Landis et al., 2021). Based on published ancestral biome reconstructions, these derived biomes were set as foreground clades, with warm temperate forests set as the background (Landis et al., 2021). Likelihood ratio tests comparing these models to their respective null models are shown in Table 2.

**Table 2.** Likelihood ratio tests of PAML clade models C and D (CmC, CmD) in *Viburnum* compared to their null models M2a_rel and M3 (Yang, 2007; Weadick and Chang, 2012). *ω*_0_–*ω*_2_ values are shown representing *d*_N_/*d*_S_ values for each site class, along with proportion of sites for each site class. In M2a_rel, *ω*_2_ is shared across all clades, whereas in CmC and CmD, *ω*_2_ is estimated separately for each class of clade. n.p., number of parameters; AIC, Akaike Information Criterion; ln*L*, log-likelihood; κ, transition/transversion ratio; *χ*^2^, chi-squared; d.f., degrees of freedom; *p*, *p*-value of likelihood ratio test. See Table 1 for M3 parameters in *Viburnum*.

| Model | n.p. | AIC | $\ln L$ | $\kappa$ | $\omega_0$ | $\omega_1$ | $\omega_2$ | $\chi^2$ | d.f. | $p$ |
| --- | --- | --- | --- | --- | --- | --- | --- | --- | --- | --- |
| M2a_rel | 298 | 7241 | -3322.5 | 1.06 | 0 (87.9%) | 1 (7.5%) | 18.9 (4.6%) |  |  |  |
| CmC: warm temperate (background) | 301 | 7236 | -3317.1 | 1.06 | 0 (88.1%) | 1 (7.2%) | 16.9 (4.8%) | 10.8 | 3 | 0.013 |
| cold temperate clades |  |  |  |  |  |  | 19.9 |  |  |  |
| tropical clades |  |  |  |  |  |  | 9.2 |  |  |  |
| cloud forest clades |  |  |  |  |  |  | 161.1 |  |  |  |
| CmD: warm temperate (background) | 302 | 7234 | -3314.8 | 1.05 | 0 (89.7%) | 1.5 (5.7%) | 16.9 (4.6%) | -7.5 | 3 | 1.00 |
| cold temperate clades |  |  |  |  |  |  | 21.2 |  |  |  |
| tropical clades |  |  |  |  |  |  | 9.9 |  |  |  |
| cloud forest clades |  |  |  |  |  |  | 158.8 |  |  |  |

CmC fit significantly better than the null model M2a_rel (*p* = 0.0129), indicating divergence in the strength of selection between biomes (Table 2). All clades were under positive selection as indicated by a class of sites *d*_N_/*d*_S_ > 1 (Table 2). Clades in cold temperate and cloud forests showed stronger positive selection than those in warm temperate forests, with *d*_N_/*d*_S_ values for the freely varying site class (*ω*_2_) of 20 and 161, respectively, compared to 17 in warm temperate forests (Table 2). In contrast, positive selection was relaxed in tropical clades, which had a lower *ω*_2_ of about 9 (Table 2). CmD, which allows for all site classes to vary freely, did not fit the data better compared to its null model M3 (*p* = 1.00), but showed a similar trend in *ω*_2_ values (Table ^2^).

To further examine this association, binary tests for correlated evolution (Pagel, 1994) were used to evaluate the dependence of transition rates of amino acid on biome and biome on amino acid at each site under positive selection or convergent evolution. Models in which amino acid and biome were interdependent had poor model fit for all sites (data not shown). Likelihood ratio tests indicated that 12 substitutions in viburnums were associated with biome transitions (Table S 4). Of the twelve amino acid substitutions that exhibited significant correlations with biome, eight showed AIC values supporting dependency of biome on amino acid. It was therefore predicted that changes in protein evolution had a greater impact on biome shifts than the reverse (Table S 4). Four substitutions, S76→N, V157→S, V255→I, and A328→S, were associated with faster transitions into warmer environments (Table S 4). M309→I was associated with faster transitions into colder environments. Stochastic character mapping predicted a greater number and rate of independent transitions into warmer environments in those derived amino acids (Figure 3A,C,H,K). In contrast, the M309→I substitution was almost always associated with cold biomes (Figure 3J). The remaining seven substitutions had derived amino acids that exhibited symmetrical transition rates (Figure 3B,D,E,F,G,I,L; Table S 4). Together, These findings suggest that substitutions in RbcL play a role in biome transitions in viburnum.

**Figure 3.**
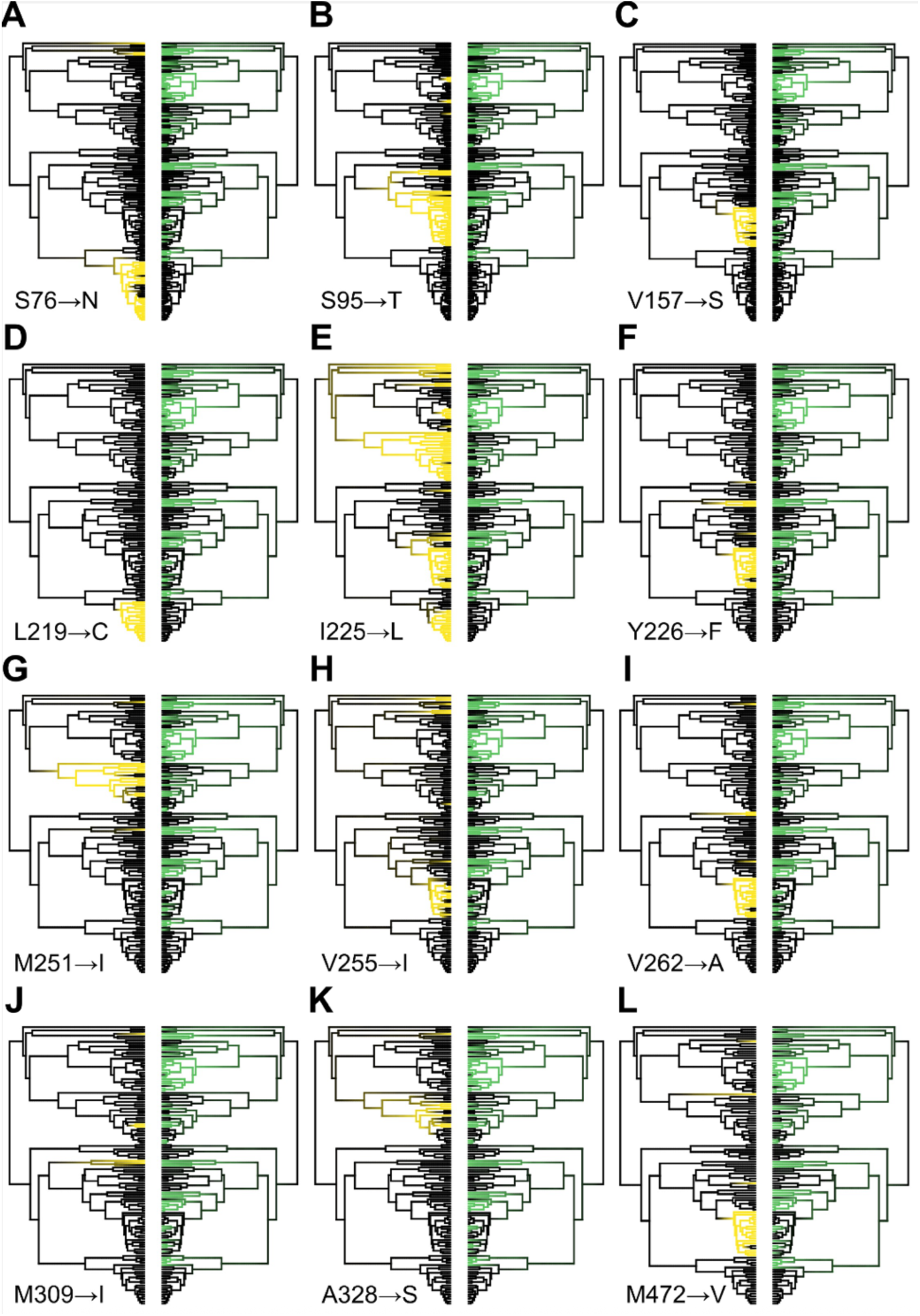
Stochastic character mapping of correlated amino acid substitutions and biome shifts in Viburnum in *Viburnum* based on Pagel (1994) tests. The best ‘fitMk’ model in ‘phytools’ was used in 100 stochastic mapping simulations. The reconstructed ancestral state for the amino acid from ‘PAML’ and biome from Landis et al. (2021) was fixed at the root. Black, ancestral amino acid or warm temperate, cloud, or tropical forests; yellow, derived amino acid; green, cold temperate forests.

### Amino acid substitutions influenced non-covalent interactions

To determine whether amino acid substitutions at positively selected or convergent sites influenced non-covalent interactions, we conducted homology modeling using a spinach crystal structure (Andersson, 1996) to assess potential impacts on protein rigidity. Eleven amino acid substitutions involved shifts towards increased ability to form hydrogen bonds (Figure 4), whereas three substitutions resulted decreased hydrogen bonding ability (Figure 5). Possible hydrogen bonding interactions were examined with other side chains within 3.7Å, a distance that encompasses weak, moderate, and strong hydrogen bonds (Jeffrey, 1997 p. 12).

**Figure 4.**
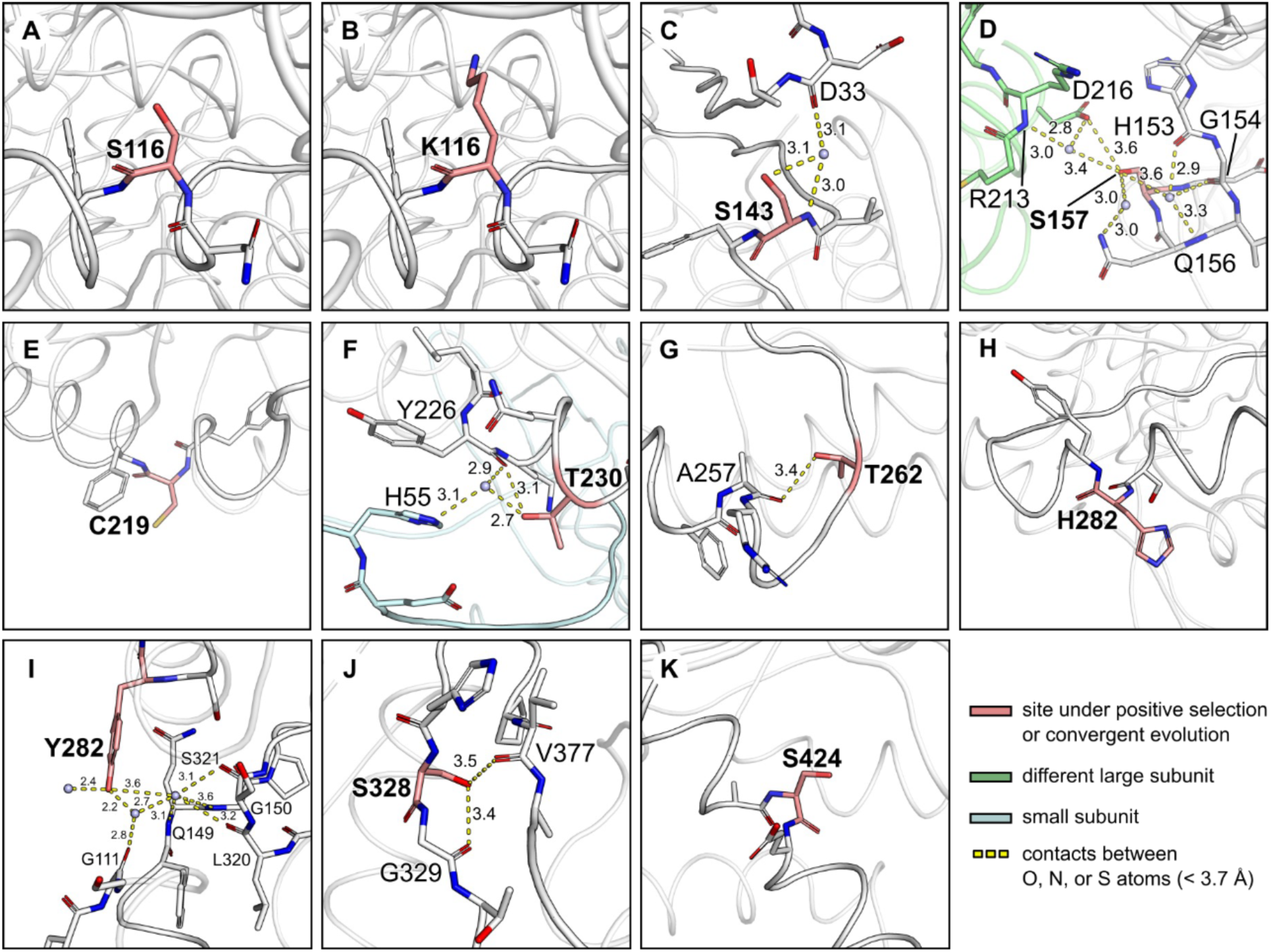
Non-covalent interactions introduced by amino acid substitutions in *Dryopteris*, *Limonium*, *Pinus*, and *Viburnum*, visualized on a spinach homology structure of the rubisco large subunit (PDB: 8RUC) (Andersson, 1996). The derived amino acid (bolded) is shown for sites that ancestrally do not contain O, N, or S atoms. Grey spheres, the position of water molecules; numbers next to dashed yellow lines, distance in Å between O, N, or S atoms.

**Figure 5.**
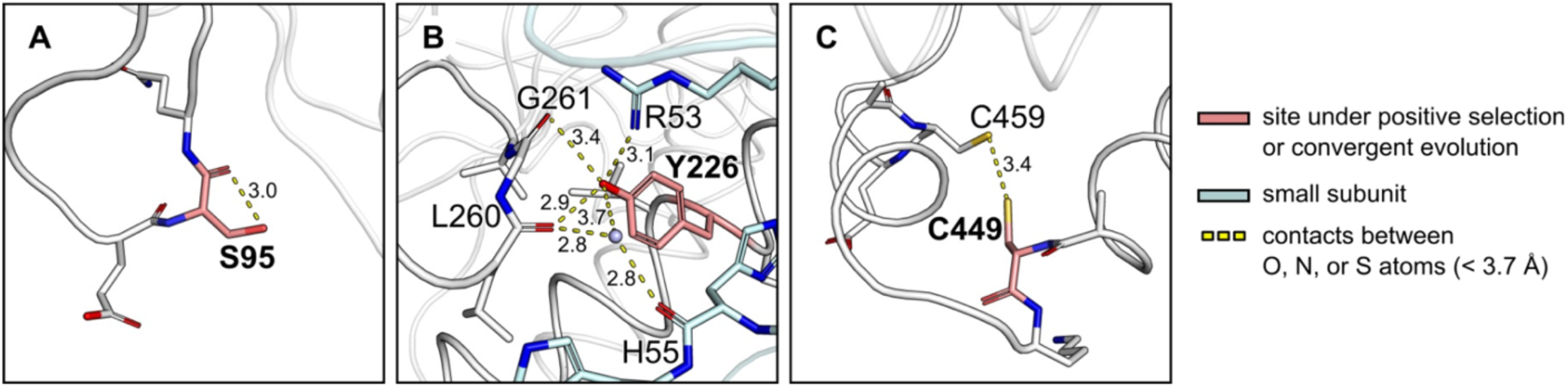
Non-covalent interactions removed by amino acid substitutions in *Dryopteris*, *Limonium*, *Pinus*, and *Viburnum*, visualized on a spinach homology structure of the rubisco large subunit (PDB: 8RUC) (Andersson, 1996). The ancestral amino acid (bolded) is shown for sites where the derived amino acid does not contain O, N, or S atoms. Grey spheres, the position of water molecules; numbers next to dashed yellow lines, distance in Å between O, N, or S atoms.

Most amino acid substitutions that increased hydrogen bonding capability involved the replacing non-polar aliphatic or aromatic residues for residues containing hydroxyl groups. About half of these replacements (S143, S157, T230, T262, Y282, and S328) were predicted to form new non-covalent interactions within the same subunit, with another large or small subunit, or with water molecules (Figure 4C,D,F,G,I,J). In contrast, non-covalent interactions were not predicted to be affected by other substitutions for polar residues (see Figure 4A,B,E,H,K). Conversely, substitutions of serine, tyrosine, or cysteine residues (S95, Y226, and C449) for non-polar aliphatic or aromatic residues were predicted to remove non-covalent interactions (Figure 5). Multiple water molecules were predicted to be involved in networks of non-covalent interactions (Figure 4C,D,F,I; Figure 5B), indicating that amino acid substitutions may alter solvent interactions by adding or removing water bridges.

### Amino acid substitutions were associated with protein stability changes

To quantify the extent to which each amino acid substitution affects holoenzyme stability, we modelled the ΔΔ*G*_fold_, an index of stability and ease of protein folding. The change in ΔΔ*G*_fold_ from the ancestral to derived amino acid in each substitution is shown in Figure 6. Derived residues that add oxygen, nitrogen, or sulfur atoms were predicted to increase stability contributions from hydrogen bonds by about −0.74 kcal mol^-1^ per large subunit (Figure 6A), by locally stabilizing specific chains. Overall, the most substantial effect of substitutions was on hydrophobic group interactions with the solvent, with a predicted mean destabilization of +1.7 kcal mol^-1^ large subunit^-1^ (Figure 6B); the addition of polar, hydrophilic atoms may therefore perturb the exposure or burial of hydrophobic groups, thereby affecting stability. Altogether, the total ΔΔ*G*_fold_, which sums all protein interactions (see Table S 1 for the remaining contributors to stability), indicates that the addition of polar atoms resulted in a predicted mean destabilization of about +1.8 kcal mol^-1^ large subunit^-1^ for each residue (Figure 6C).

**Figure 6.**
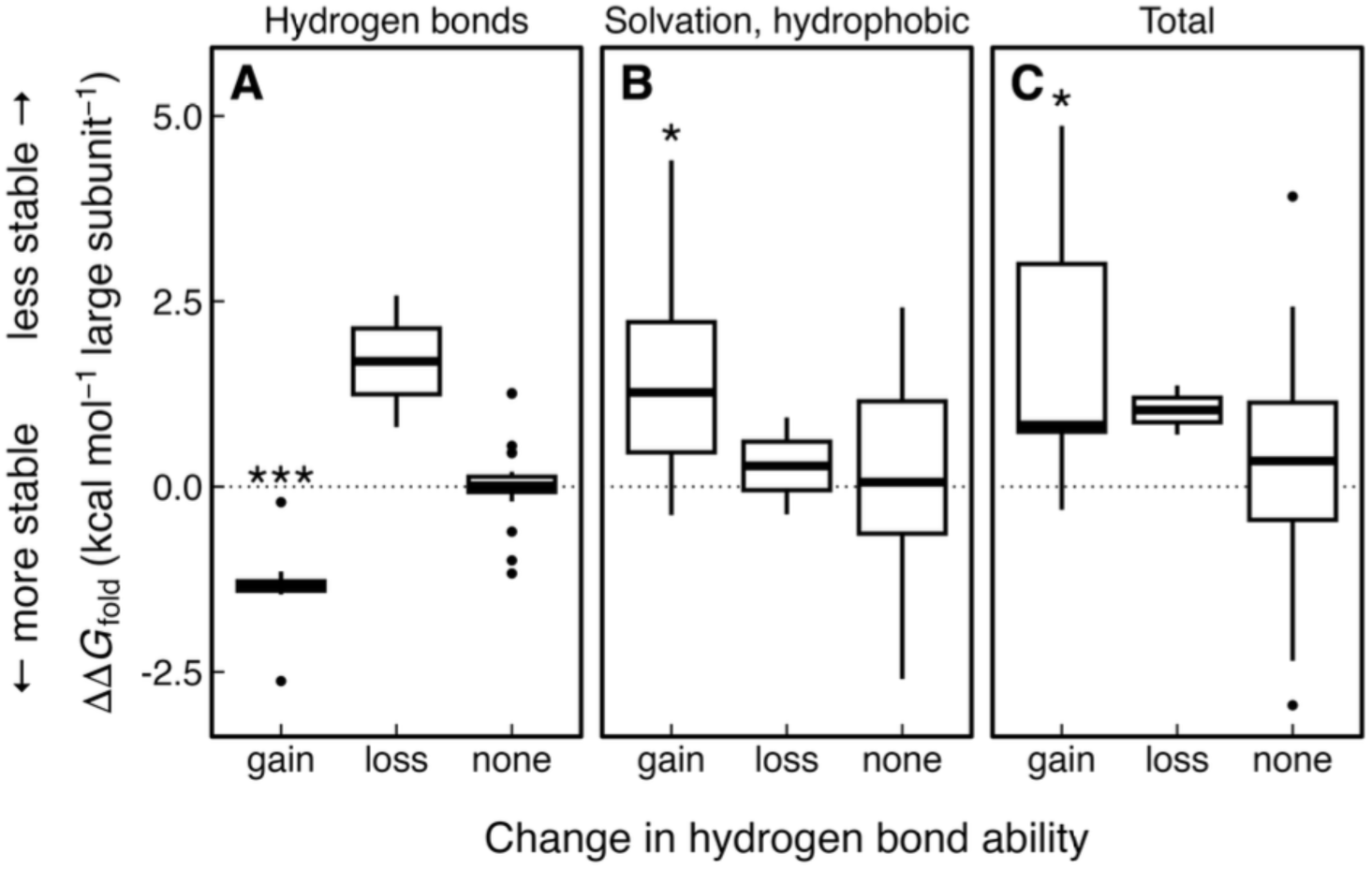
Modelled change in free energy of folding (ΔΔ*G*_fold_) of the rubisco holoenzyme from ancestral to derived amino acid in *Dryopteris*, *Limonium*, *Pinus*, and *Viburnum*. ΔΔ*G*_fold_ values were predicted for each amino acid substitution at sites under positive selection and/or convergent evolution. ΔΔ*G*_fold_ values are grouped by: ancestral amino acid side chain unable to form hydrogen bonds and derived able (“gain”); both ancestral and derived either able or unable (“none”); and ancestral able and derived unable to form hydrogen bonds (“loss”). Boxplots show the distribution of ΔΔ*G*_fold_ (from three replicate simulations), standardized to each of eight large subunits in the holoenzyme. Symbols represent the mean for each category of substitution. (A) and (B), contribution from hydrogen bonds and interaction of hydrophobic groups with the solvent, respectively; (C) total ΔΔ*G*_fold_. See Table S2 for other components. One sample t-tests were used to test if each type of substitution had ΔΔ*G_f_*_old_ values different from zero; *, *p* < 0.05; **, *p* < 0.01; ***, *p* < 0.001.

### Stability was weakly correlated with growing season temperature

To evaluate the additive effects of amino acid diversity on protein stability, the large subunit sequence in the spinach homology model was mutated to predict the ΔΔ*G*_fold_ of each gap-free sequence in this study. The small subunit sequences were not mutated. Across wood ferns, sea lavenders, pines, and viburnums, maximum growing season temperature had a modest but statistically significant correlation with modeled ΔΔ*G*_fold_ (*p* = 0.0013; *R*^2^ = 0.028; Figure 7A). Given the low variance explained and loss of signal after phylogenetic correction, these associations should be interpreted with caution. Climatic variables interpolated from weather station data is a coarse representation of the actual growing conditions. The most stable tercile of rubiscos occurred in environments that are 2.4 °C warmer than those of the least stable tercile. (Figure 7B). The relationship between ΔΔ*G*_fold_ and growing season temperature, when analyzed using PGLS was altered when using a binned approach (Figure 7B) and disappeared when using continuous data. Together, the analyses indicate a strong phylogenetic effect in rubisco thermal adaptation.

**Figure 7.**
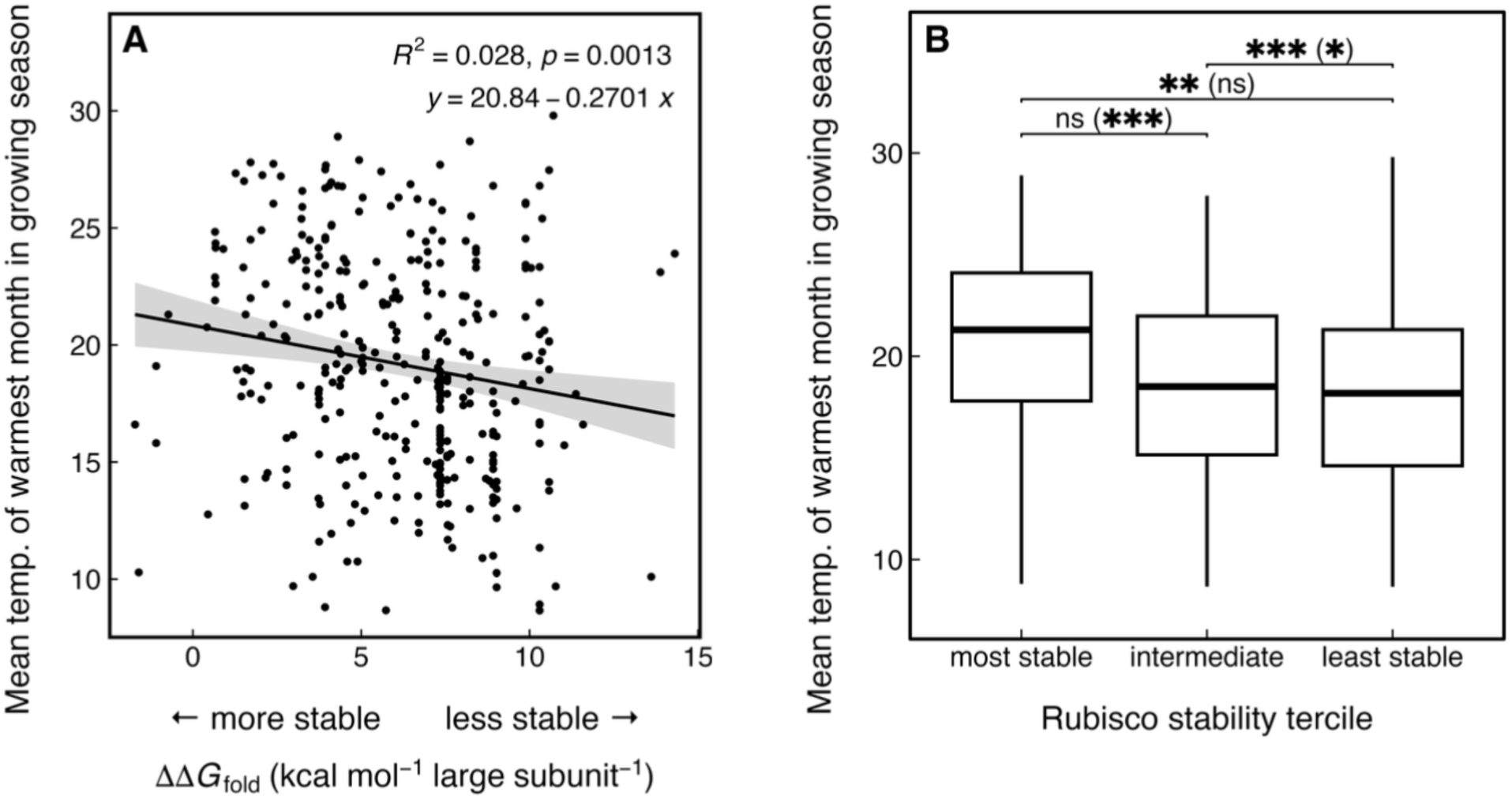
(A) Correlations between mean temperature of the warmest month in the growing season and change in free energy of folding (ΔΔ*G*_fold_) of rubisco. The solid line is a linear regression, and the grey shaded area represents the 95% confidence interval. This correlation is not significant using phylogenetic generalized least squares (PGLS). Holoenzyme ΔΔ*G*_fold_ values were predicted with each gap-free *rbcL* sequence (*n* = 399 species with 174 unique sequences) by mutating the spinach crystal structure (PDB: 8RUC). The mean values from three replicate runs in ‘FoldX’ are shown. The growing season was defined as the months with monthly mean temperature ≥ 5°C and aridity index > 0.05 (Wright et al., 2017; He et al., 2020). See Figure S 1 for regressions on each genus. (B) ΔΔ*G*_fold_ values were also analyzed with a binning approach, which is standard practice (Studer et al. 2014). Each tercile is 33% of the dataset. Boxplots show the distribution of mean temperature of the warmest month in the growing season from at least three GBIF occurrences. Asterisks indicate a significant difference based on *t*-tests on log-transformed data. Asterisks in parentheses indicate a significant difference based on PGLS analyses. *, *p* < 0.05; **, *p* < 0.01; ***, *p* < 0.001

## Discussion

### Convergent modification of the rubisco large subunit gene

Focusing on *rbcL*, we replicated analyses in wood ferns, sea lavenders, pines, and viburnums and found pervasive positive selection and/or convergent evolution, suggesting that adaptive evolution occurred (Table 1; Table 2). A small fraction of sites (95, 226, 251, 255, 262, 340, 375, and 449) were consistently under positive selection in multiple genera (Table 1; Figure 2).

However, the disparate patterns of sites targeted by positive selection highlight the lineage-specific effects and the significance of evolutionary history in the evolution of rubisco (Bouvier et al., 2021). Viburnums accounted for over 40% of those amino acid sites identified in this study, which may reflect its repeated shifts into tropical and cold temperate biomes (Table S 4; Landis et al., 2021). Moreover, although tests of positive selection are routinely performed for *rbcL*, another indicator of adaptive evolution is the occurrence convergent amino acid substitutions on multiple branches of a phylogeny. Convergent or parallel evolution in rubisco has been shown previously by identifying multiple branches with the same derived amino acid (Kapralov and Filatov, 2007; Christin et al., 2008). However, introgression through plastid capture can be a confounding factor in *rbcL* (Yao et al., 2019). We used a recently developed method to account for these errors (Fukushima and Pollock, 2023) and show that a high level of convergence is not due to introgression. These results support the hypothesis that *rbcL* has been repeatedly modified in a manner that would indicate adaptive evolution.

### Interactions with the solvent, not hydrogen bonds, best explain protein stability changes

Unlike LDH or MDH, rubisco is constrained by dual substrate specificity, such that stability, turnover rate, and CO_2_ specificity must be fine-tuned. Structural changes can be subtle; a five-fold difference in *S*_c/o_ can result from a free energy difference smaller than that of a hydrogen bond (Spreitzer and Salvucci, 2002). In lineages examined here, 11 and 3 unique substitutions added and removed oxygen, nitrogen, or sulfur atoms in the amino acid side chains, respectively, which we interpreted as a change in hydrogen bond ability (Figure 4; Figure 5). Six and three of those substitutions were predicted to add or remove interactions within 3.7 Å, respectively (Figure 4A,B,E,H,K; Figure 5). These interactions are hypothesized to be hydrogen bonds, water bridges, or disulfide bridges, which would stabilize the protein. An increase in polarity is generally hypothesized to stabilize metabolic enzymes in poikilotherms through hydrogen bonds, which typically increase stability by 3–9 kcal mol^-1^ (Fersht, 1995; Dong and Somero, 2009). In contrast to this hypothesis, the increases in polarity were predicted to decreased holoenzyme stability on average by 1.8 kcal mol^-1^ per large subunit (Figure 6C). Interactions of hydrophobic residues with the solvent appeared to destabilize the rubisco holoenzyme moderately (Figure 6B). This pattern diverges from the patterns observed in fish and mollusc LDH and MDH (Dong and Somero, 2009; Fields et al., 2015). We propose that it is essential to consider the stabilizing effect of water bridges and especially those involving waters buried in the protein (Park and Saven, 2005). It is also worth noting that all modelling was performed while keeping RbcS unchanged to isolate the effects of amino acid substitutions in RbcL. An alternative explanation is, therefore, that changes to substitutions in RbcL is coordinated with RbcS. We suggest further investigation.

### Rubisco large subunit evolution is associated with ecology and ecological shifts

Divergence in positive selection between species in ancestral and derived biomes was tested in viburnums using previously published biome scorings (Landis et al., 2021). Clade model analyses with CmC revealed that transitions to cold temperate and cloud forest clades were associated with stronger positive selection, while tropical clades experienced weakened positive selection (Table 2). This is interesting because transitions into cold temperate forests in *Viburnum* often involve the evolution of deciduousness (Edwards et al., 2017), buffering differences in leaf environment. Additionally, the cloud forest clade has a strikingly high *d*_N_/*d*_S_ value of 161, which may reflect the unique, island-like nature of its disjunct cloud forest habitats (Donoghue et al., 2022). CmD showed similar trends but did not fit the data better than its null.

Additionally, the correlation of amino acid transition rates with those of biomes was assessed using phylogenetic comparative methods (Pagel, 1994). Focusing on transition rates rather than simply correlating traits allows for inferences about trait dependencies (Pagel, 1994 p. 199).

While protein modifications have been linked to ecological shifts in animal systems (Dong and Somero, 2009; Schott et al., 2014; Dungan et al., 2016), fewer studies have done so for rubisco (but see Hermida-Carrera et al., 2017). A large fraction of amino acid shifts in viburnums were associated with symmetrical transition rates, implying no strong preference for a particular biome (Table S 4). In the substitutions that do correlate with biome shifts, the analyses suggest that RbcL evolution can either facilitate or inhibit transitions into new habitats, but that these shifts are not necessary or rapid (Figure 3). The data therefore suggest a role of rubisco evolution in facilitating habitat shifts in “replicated bursts” (Maddison and FitzJohn, 2015), which is when one trait is sufficient but not necessary for influencing the evolution of another trait. In short, the data demonstrate that RbcL evolution influenced biome shifts in *Viburnum* but did not fully explain them.

Using an estimation of the growing season temperature (see Materials and Methods), we found that rubiscos with higher predicted stability occurred in warmer climates. This relationship parallels documented relationships in Δ*G* of unfolding with melting temperature, which in turn correlates with the temperature of maximal stability (Rees and Robertson, 2001). Although we did not collect rubisco kinetics data, our combined analyses support a hypothesis that proteins from different species are fine-tuned for their environment, such that they perform similarly in their respective operating conditions. The weakened correlations when using PGLS for comparisons highlights the role of phylogenetic constraint in rubisco evolution (Bouvier et al., 2021). The 2.4 °C difference between the most and least stable terciles would correspond to a 21% difference in rubisco activity with a Q_10_ of 2.2 (Cen and Sage, 2005).

### Rubisco, rubisco activase, and chaperones as an evolutionary module: phylogenetic considerations

What might explain phylogenetic patterns in amino acid substitutions, holoenzyme stability, and climate? As argued by (Bouvier et al., 2021), complementarity between rubisco subunits, rubisco activase, and chaperones could influence rubisco evolution, a pattern we anticipate would manifest as part of the phylogenetic signal. We suggest that rubisco subunits and related proteins act as an evolutionary module: a semi-independent set of traits that coevolve due to functional integration. In modular systems, traits evolve together because they contribute to a shared function, yet remain relatively decoupled from traits outside the module (Wagner, 1996). For rubisco, although catalytic activity is localized to the large subunit, its folding, assembly, activation, and overall function are dependent on interactions with the small subunit and a suite of chaperones and auxiliary factors. Modular organization can arise because characters contributing to organismal function often evolve in coordination. This functional interdependence likely constrains evolutionary trajectories, to varying degrees, producing coordinated shifts in protein-protein and protein-solvent interactions across lineages.

Evidence from recombinant expression supports this view that rubisco and related proteins function as a module. The chaperones Raf1, Raf2, RbcX, BSD2, and the Cpn60/20 chaperonin complex are required for a functional holoenzyme, defining a minimal set of proteins required for assembly. RbcL and RbcS do not assemble without chaperones, although expressing the cognate Raf1 partially rescues assembly (Whitney et al., 2015; Aigner et al., 2017). Beyond assembly, impairment of rubisco activase expression decreases the CO_2_ assimilation rate, including in non-steady state conditions (Jiang et al., 1994; Hammond et al., 1998; Taylor et al., 2022), and the expression of specific rubisco activase orthologs are required for proper function of rubisco (Milward, 2018; Gunn et al., 2020). Within the holoenzyme, the small subunit and its isoforms can have a considerable effect on rubisco activity in variable environments, further underscoring the enzyme’s sensitivity to its broader molecular context (Wostrikoff and Stern, 2007; Ishikawa et al., 2011; Lin et al., 2020; Cavanagh et al., 2022).

Patterns of protein integration are not unique to rubisco; in viral proteins, for instance, host chaperones can constrain the regions of the protein that is evolutionarily accessible, thereby shaping its evolutionary trajectory (Yoon et al., 2022). Such constraints likely reflect the degree of integration among molecular components. Importantly, the strength of modular integration is itself variable, and this variation can influence how modules evolve over time (Wagner, 1996; Wagner et al., 2007). The presence of multiple isoforms (and differing copy numbers) of RbcS and activase (Salvucci et al., 1987; Spreitzer, 2003) suggests that integration is variable in rubisco, such that RbcL may not always reflect modifications other interacting enzymes. Deep-time divergences in rubisco and likely elements beyond RbcL may be linked to deviations from the canonical *k*_cat_^c^ and *K*_c_ trade-off when looking at large data sets (Flamholz et al., 2019), such as in hornworts vs. other land plants (Oh et al., 2024). Given the conformational constraints that RbcS imposes on RbcL (Amritkar et al., 2025), it is worth considering how RbcS could reciprocally influence RbcL. This highlights how the functional context of RbcL has evolved, such that relying solely on RbcL to interpret kinetic variation may overlook key co-evolutionary shifts (Bouvier et al., 2021). As a result, evolutionary history within the green lineage can obscure the drivers of rubisco adaptation if other components of the module are not considered. We suggest that treating rubisco and its interacting partners as an evolutionary module offers a valuable framework for understanding rubisco evolution.

To better examine rubisco evolution in nature, a model clade approach may minimize confounding effects of deep phylogenetic divergences (Donoghue and Edwards, 2019; Mabry et al., 2024). This strategy is valuable because photosynthesis is the outcome of nested processes that each can become rate-limiting (Farquhar et al., 1980). As such, the rubisco module cannot be meaningfully removed from the broader organismal context; factors such as photochemistry, anatomy, hydraulics, and phenology affect the operating conditions of rubisco and in turn reflect the evolutionary history of plant lineages. Thus, dense and comprehensive sampling in a phylogenetic context (e.g., small subunit, activase, and chaperones) and the temperature response of rubisco kinetics in a well-studied model clade should provide a better picture of enablers and inhibitors of rubisco evolution. Focusing on lineages that have independently shifted between climatic regimes allows for evolutionary comparisons of the thermal adaptations. For instance, the *Dentata*, *Mollotinus*, and *Urceolata* clades within *Viburnum* are groups that each transitioned from warm to cold environments in the past 15 Ma (Landis et al., 2021) and may be suitable for characterization of rubisco kinetics and its thermal responses (Orr et al., 2016; Sharwood et al., 2016; Sargent et al., 2025).

### Conclusion

Our analyses across lineages indicate that patterns in RbcL evolution resemble those seen in enzymes from thermally diverse animals, suggesting shared constraints and selective pressures; however, the complexity of the rubisco system may impose deep phylogenetic constraints.

Positive selection, convergent evolution, altered side chain interactions in the homology model, correlation of *d*_N_/*d*_S_ and amino acid substitutions with biome in *Viburnum*, and weak correlation of stability with growing season temperature together suggest a role of rubisco in thermal adaptation. However, the correlation between protein stability and temperature was weakened after correcting for phylogeny, underscoring the influence of evolutionary history on these patterns. To disentangle environmental effects from phylogenetic effects, we recommend a model clade approach that enables focused comparisons within shared phylogenetic backgrounds. A more phylogenetically grounded understanding of rubisco in natural systems could then dovetail into efforts to predict and improve plant resilience.

## Data availability

All data and code required to reproduce the analyses are deposited on GitHub (github.com/aleungplants/rubisco-evolution).

## Author contributions

A.L. performed the research with guidance from B.S.W. and R.F.S.

## Acknowledgements

The authors thank Drs. Emily Sessa and Konstantina Koutroumpa for providing the *Dryopteris* and *Limonium* tree files and giving permission to provide these files in the supplemental data. The research was funded by a Queen Elizabeth II/Charles E. Eckenwalder Scholarship in Science and Technology to A.L. and NSERC Discovery Grants to B.S.W.C. and R.F.S.

## Supplementary material

**Table S 1.** Results of substitution saturation assessments in DAMBE v7 (Xia, 2018). Observed indices of substitution saturation (*I*_ss_) are compared to critical values for symmetrical (*I*_ss.cSym_) and asymmetrical (*I*_ss.cAsym_) topologies. *t*, *df*, and *p* report the t-statistic, degrees of freedom, and significance for testing whether *I*_ss_ is significantly lower than *I*_ss.c_. A significant *I*_ss_ < *I*_ss.c_ indicates little saturation.

| Genus | Number of taxa | $I_{ss}$ | $I_{ss,cSym}$ | $t$ | $df$ | $p$ | $I_{ss,cAsym}$ | $t$ | $df$ | $p$ |
| --- | --- | --- | --- | --- | --- | --- | --- | --- | --- | --- |
| *Dryopteris* | 4 | 0.180 | 0.828 | 45.133 | 603 | <0.0001 | 0.796 | 42.931 | 603 | <0.0001 |
|  | 8 | 0.202 | 0.800 | 39.577 | 603 | <0.0001 | 0.697 | 32.780 | 603 | <0.0001 |
|  | 16 | 0.204 | 0.784 | 38.195 | 603 | <0.0001 | 0.594 | 25.671 | 603 | <0.0001 |
|  | 32 | 0.212 | 0.763 | 32.096 | 603 | <0.0001 | 0.471 | 15.065 | 603 | <0.0001 |
| *Limonium* | 4 | 0.909 | 0.828 | 5.274 | 198 | <0.0001 | 0.797 | 7.328 | 198 | <0.0001 |
|  | 8 | 0.959 | 0.801 | 8.721 | 198 | <0.0001 | 0.698 | 14.363 | 198 | <0.0001 |
|  | 16 | 1.034 | 0.784 | 12.542 | 198 | <0.0001 | 0.595 | 22.071 | 198 | <0.0001 |
|  | 32 | 1.269 | 0.764 | 22.509 | 198 | <0.0001 | 0.472 | 35.537 | 198 | <0.0001 |
| *Pinus* | 4 | 0.040 | 0.824 | 76.304 | 401 | <0.0001 | 0.792 | 73.195 | 401 | <0.0001 |
|  | 8 | 0.038 | 0.794 | 69.751 | 401 | <0.0001 | 0.690 | 60.138 | 401 | <0.0001 |
|  | 16 | 0.038 | 0.778 | 67.975 | 401 | <0.0001 | 0.584 | 50.184 | 401 | <0.0001 |
|  | 32 | 0.039 | 0.756 | 65.060 | 401 | <0.0001 | 0.457 | 37.973 | 401 | <0.0001 |
| *Viburnum* | 4 | 0.429 | 0.830 | 47.869 | 460 | <0.0001 | 0.799 | 44.139 | 460 | <0.0001 |
|  | 8 | 0.459 | 0.803 | 37.779 | 460 | <0.0001 | 0.701 | 26.588 | 460 | <0.0001 |
|  | 16 | 0.496 | 0.787 | 30.640 | 460 | <0.0001 | 0.599 | 10.845 | 460 | <0.0001 |
|  | 32 | 0.559 | 0.767 | 21.077 | 460 | <0.0001 | 0.478 | 8.236 | 460 | <0.0001 |

**Table S 2.** Change in free energy of folding (ΔΔ*G*) of the rubisco holoenzyme for each amino acid substitutions at sites under positive selection and/or convergent evolution. ΔΔ*G* values are the mean from ten simulations in FoldX 5.1 for the whole holoenzyme. Contributions from hydrogen bonds shown in Fig. 5 were summed from the backbone and sidechain hydrogen bond contributions. SD, standard deviation of the total ΔΔ*G* from ten simulations.

| Contributions to holoenzyme $\Delta\Delta G$ (kcal mol<sup>-1</sup>) | | | | | | | | | | | | | |
| --- | --- | --- | --- | --- | --- | --- | --- | --- | --- | --- | --- | --- | --- |
| Genus | Site | Substitution | Total $\Delta\Delta G$<br>(kcal mol<sup>-1</sup>) | SD | H-bonds,<br>backbone | H-bonds,<br>sidechain | Solvation,<br>polar | Solvation,<br>hydrophobic | Clashes,<br>VanderWaals | Clashes,<br>torsional | Clashes,<br>backbone | VanderWaals | Electrostatics |
| *Dryopteris* | 116 | L116K | 25.81 | 0.46 | -0.36 | -4.14 | 19.37 | 2.95 | -0.17 | 0.06 | 0.16 | -1.71 | 2.34 |
|  |  | L116M | 2.63 | 0.13 | -0.12 | 0.11 | -0.09 | -2.82 | -0.32 | -0.14 | 0.01 | -2.74 | 0.00 |
|  |  | L116P | 57.44 | 0.52 | 8.52 | 0.16 | -0.14 | 15.60 | 36.86 | -0.23 | -18.17 | 6.95 | 0.32 |
|  |  | L116S | 33.25 | 0.22 | -1.97 | -7.20 | 4.04 | 28.78 | -0.05 | -0.31 | -0.19 | 11.25 | 0.00 |
|  | 262 | A262V | -16.12 | 0.10 | -0.87 | -0.01 | 6.67 | -18.67 | 0.01 | 0.04 | 4.02 | -10.18 | 0.12 |
|  | 282 | F282H | 6.02 | 0.05 | -5.42 | -7.11 | 6.83 | 6.45 | -0.17 | -0.81 | -0.24 | 2.22 | -0.88 |
|  |  | F282I | 5.64 | 0.36 | 0.05 | -0.55 | 0.72 | 4.09 | 0.24 | -0.56 | 0.98 | 2.80 | 0.02 |
|  |  | F282L | 0.84 | 0.44 | -0.29 | 0.06 | -0.99 | 2.50 | 0.04 | -0.73 | 0.25 | 2.41 | 0.14 |
|  |  | F282Y | 2.20 | 0.09 | 0.00 | -0.09 | 3.12 | 0.12 | 0.00 | -0.46 | 0.00 | -0.81 | -0.02 |
|  | 375 | L375F | 21.79 | 4.77 | -0.34 | 3.26 | 5.90 | -1.53 | 26.09 | 1.56 | 2.07 | -4.78 | -5.06 |
|  |  | L375I | 4.00 | 0.73 | -0.30 | -0.22 | 0.20 | -0.40 | 1.51 | 4.73 | 0.13 | -0.13 | 0.90 |
|  | 424 | L424S | 40.46 | 0.02 | -2.65 | -6.55 | 3.05 | 33.97 | 0.57 | -0.24 | -0.43 | 13.90 | 0.31 |
| *Limonium* | 43 | T43S | -2.75 | 1.06 | 1.85 | -3.87 | -2.96 | 8.18 | -3.80 | -6.88 | -0.55 | 4.00 | 0.01 |
|  | 95 | N95S | 17.05 | 0.28 | 0.74 | 10.00 | -7.01 | 5.75 | 0.72 | -0.97 | -0.55 | 5.33 | 0.06 |
|  | 245 | G245A | 8.92 | 0.39 | 1.07 | -0.82 | 7.38 | -7.20 | 15.66 | -0.04 | 2.25 | -5.20 | -0.24 |
|  | 309 | I309M | 2.68 | 0.29 | -0.01 | -0.01 | -0.57 | -4.11 | 0.02 | 0.17 | -0.48 | -3.98 | 0.15 |
|  | 340 | D340E | -4.35 | 0.19 | 0.21 | 0.19 | -1.12 | -1.96 | 0.06 | -0.01 | 0.52 | -1.04 | -1.83 |
| Genus | Site | Substitution | Total $\Delta\Delta G$<br>(kcal mol<sup>-1</sup>) | SD | H-bonds,<br>backbone | H-bonds,<br>sidechain | Solvation,<br>polar | Solvation,<br>hydrophobic | Clashes,<br>VanderWaals | Clashes,<br>torsional | Clashes,<br>backbone | VanderWaals | Electrostatics |
| *Pinus* | 4 | K4Q | ND | ND | ND | ND | ND | ND | ND | ND | ND | ND | ND |
|  | 50 | P50A | 15.91 | 0.04 | 0.51 | 0.00 | 0.62 | 3.88 | -0.01 | 0.06 | 7.36 | 2.33 | -0.01 |
|  | 95 | S95A | -0.93 | 1.03 | 2.36 | 2.97 | -2.35 | -1.34 | -0.17 | -0.15 | -0.03 | 0.50 | 0.03 |
|  | 95 | S95N | -16.96 | 0.67 | -0.69 | -9.61 | 7.31 | -5.84 | -1.29 | 1.07 | 0.58 | -5.49 | -0.06 |
|  | 95 | S95T | 4.80 | 0.91 | 2.33 | 2.97 | 1.22 | -3.86 | -0.96 | 2.43 | 0.43 | -1.92 | -0.02 |
|  | 143 | A143S | 8.95 | 0.13 | -1.97 | -2.29 | 6.27 | 2.84 | 0.18 | -0.06 | 0.92 | -2.25 | -0.01 |
|  | 226 | N226Y | -36.25 | 0.65 | -1.81 | -3.56 | 5.32 | -15.12 | -0.81 | -0.69 | -1.15 | -8.71 | -18.23 |
|  | 251 | M251I | 9.18 | 0.18 | 0.24 | -0.08 | 0.03 | 9.55 | 0.02 | 0.48 | -0.07 | 5.59 | -0.03 |
|  |  | M251V | 19.68 | 0.11 | 0.75 | -1.32 | -2.20 | 20.74 | 0.06 | 0.67 | -1.33 | 11.54 | -0.31 |
|  | 254 | A254V | -5.76 | 2.51 | -1.56 | 1.61 | 0.59 | -16.94 | 11.94 | 0.34 | 1.14 | -8.03 | 0.00 |
|  | 255 | V255I | -4.11 | 0.14 | -0.66 | 0.11 | 1.74 | -5.62 | 0.53 | 0.64 | 1.12 | -3.16 | 0.04 |
|  | 265 | V265I | -6.93 | 0.07 | -0.02 | 0.00 | -0.14 | -9.42 | 0.11 | -0.03 | 0.19 | -4.22 | 0.00 |
|  | 375 | L375M | 3.35 | 0.12 | -0.07 | -0.21 | 2.25 | -2.99 | -0.03 | -0.06 | 0.35 | -4.04 | -0.78 |
|  | 418 | V418A | 15.09 | 0.47 | 0.72 | 0.26 | -4.39 | 15.51 | 0.58 | -0.14 | -1.05 | 8.41 | -0.24 |
|  | 449 | C449A | 4.48 | 0.35 | 1.96 | 8.83 | -3.28 | 7.39 | -0.06 | -0.17 | -1.87 | 6.52 | 0.05 |
|  |  | C449S | 3.35 | 0.16 | -3.25 | -2.16 | 0.52 | 10.95 | -0.05 | -0.12 | -0.64 | 5.64 | -0.01 |
|  | 471 | V471A | -0.71 | 0.66 | 0.43 | 0.57 | -9.77 | 11.94 | -4.71 | -2.20 | -1.39 | 7.94 | 0.86 |
| Genus | Site | Substitution | Total $\Delta\Delta G$<br>(kcal mol<sup>-1</sup>) | SD | H-bonds,<br>backbone | H-bonds,<br>sidechain | Solvation,<br>polar | Solvation,<br>hydrophobic | Clashes,<br>VanderWaals | Clashes,<br>torsional | Clashes,<br>backbone | VanderWaals | Electrostatics |
| *Viburnum* | 76 | S76N | -8.81 | 1.28 | 2.19 | -3.90 | 4.48 | -6.36 | -1.22 | 1.29 | 2.58 | -3.63 | -0.03 |
|  | 95 | S95T | 4.85 | 0.89 | 2.10 | 3.11 | 1.15 | -3.83 | -0.63 | 2.32 | 0.41 | -1.89 | -0.03 |
|  | 157 | V157S | 7.49 | 0.65 | 0.94 | -9.92 | -5.13 | 13.74 | -0.04 | 0.08 | -2.00 | 6.27 | 1.15 |
|  | 219 | L219C | 22.26 | 0.11 | 1.53 | 0.12 | -9.01 | 16.50 | -0.26 | -0.28 | -2.49 | 8.92 | 0.38 |
|  |  | L219V | 9.38 | 0.09 | 0.23 | 0.40 | -5.52 | 8.68 | 0.15 | 0.78 | -1.12 | 5.38 | 0.54 |
|  | 225 | I225L | -5.52 | 0.19 | 0.41 | 0.01 | 0.42 | -1.69 | -3.67 | 0.57 | -1.11 | -0.91 | 0.00 |
|  | 226 | Y226F | 11.47 | 0.87 | 5.46 | 10.73 | -4.19 | -1.37 | 0.14 | -0.05 | -0.04 | 1.23 | 1.23 |
|  | 230 | A230T | 0.92 | 0.05 | -4.10 | -3.59 | 4.98 | -0.55 | 0.04 | 0.22 | 1.43 | -1.67 | -0.35 |
|  | 249 | E249D | 3.11 | 0.35 | 10.34 | 2.49 | -7.27 | 4.52 | -0.62 | -0.69 | -1.49 | 4.01 | -1.72 |
|  | 251 | M251I | 9.18 | 0.13 | 0.08 | -0.18 | 0.16 | 9.49 | 0.03 | 0.60 | -0.04 | 5.56 | -0.05 |
|  | 255 | V255I | -4.12 | 0.11 | -0.67 | 0.32 | 1.66 | -5.62 | 0.56 | 0.66 | 1.16 | -3.15 | 0.11 |
|  | 262 | V262A | 16.09 | 0.08 | 0.83 | 0.03 | -6.69 | 18.68 | -0.01 | -0.05 | -4.00 | 10.19 | -0.12 |
|  |  | V262T | 11.58 | 0.09 | -2.24 | -2.18 | 2.34 | 9.87 | -0.02 | 0.17 | -0.88 | 3.09 | 0.00 |
|  | 279 | T279S | 8.77 | 0.07 | 0.50 | 1.04 | -2.53 | 7.62 | -0.32 | -1.42 | -0.87 | 3.39 | -0.06 |
|  | 309 | M309I | -2.72 | 0.24 | -0.01 | -0.01 | 0.62 | 4.12 | -0.07 | -0.07 | 0.50 | 3.93 | -0.13 |
|  | 317 | A317G | 18.47 | 0.05 | 0.70 | 0.09 | -1.89 | 10.68 | -0.03 | -0.02 | -0.74 | 5.44 | -0.04 |
|  | 328 | A328G | 8.46 | 0.02 | 0.39 | 0.00 | -3.13 | 8.41 | -0.01 | -0.03 | -0.73 | 4.53 | -0.01 |
|  |  | A328S | 1.63 | 0.01 | -2.87 | -2.64 | 5.76 | 2.53 | 0.01 | 0.20 | 0.37 | -1.91 | 0.01 |
|  | 340 | E340D | 4.35 | 0.19 | -0.21 | -0.19 | 1.12 | 1.96 | -0.06 | 0.01 | -0.52 | 1.04 | 1.83 |
|  | 449 | C449A | 4.48 | 0.35 | 1.96 | 8.83 | -3.28 | 7.39 | -0.06 | -0.17 | -1.87 | 6.52 | 0.05 |
|  |  | C449S | 3.23 | 0.12 | -3.25 | -1.64 | 0.06 | 11.12 | -0.13 | -0.11 | -0.63 | 5.88 | -0.04 |
|  | 461 | V461I | -3.46 | 0.10 | 0.05 | 0.12 | 2.43 | -5.42 | 0.00 | 0.32 | 1.38 | -3.04 | -0.04 |
|  | 472 | M472V | 2.51 | 0.24 | 0.14 | 1.08 | -4.67 | 7.86 | -0.40 | -0.09 | -0.11 | 6.80 | 0.80 |
|  | 475 | L475I | 0.81 | 0.62 | -0.11 | 0.08 | -1.19 | 2.60 | -1.12 | -0.33 | -1.26 | 1.62 | 0.13 |

**Table S 3.** Error-corrected convergent amino acid substitutions in each genus (Fukushima and Pollock, 2023). Convergent substitutions are defined as a transition from any amino acid to the same amino acid in multiple branches of the same phylogeny. Branch numbers refer to distinct branches on which each substitution in parallel (see Data Availability for tree files; Sessa et al., 2017; Jin et al., 2021; Koutroumpa et al., 2021; Landis et al., 2021).

| Genus | Site | Amino acid at<br>the root of<br>phylogeny | Substitution | Branch numbers |
| --- | --- | --- | --- | --- |
| *Dryopteris* | 116 | L | L116→S | 32, 36, 51, 55, 56, 62, 64, 79, 225, 228, 259, 266, 337 |
| · | 262 | A | A262→V | 4, 55, 62, 112, 200, 202, 203, 207, 208, 213,<br>215, 249, 275, 280, 281, 286, 289, 316, 361 |
| · | 282 | F | F282→H | 103 |
| · | · | · | F282→Y | 149, 353 |
| · | · | · | Y282→H | 32, 40, 94, 95, 282, 337, 344, 345 |
| · | 375 | L | L375→F | 2, 43, 59, 60, 151, 202, 208, 281 |
| · | 424 | · | L424→S | 2, 62, 112, 116, 225, 294, 361, 362 |
| *Limonium* | 43 | T | T43→S | 271, 355 |
| · | 95 | N | N95→S | 60, 63, 112, 247, 336 |
| · | 309 | I | I309→M | 139, 193 |
| *Pinus* | 4 | K | K4→Q | 52, 83, 85, 119, 146, 154 |
| · | 50 | P | P50→A | 139, 190 |
| · | 95 | S | S95→N | 139, 190 |
| · | · | · | S95→T | 80, 90 |
| · | 143 | A | A143→S | 14, 53 |
| · | 226 | N | N226→Y | 10, 40, 180 |
| · | 251 | M | I251→V | 13, 27 |
| · | · | · | M251→I | 10, 53, 79, 127, 180 |
| · | 254 | A | A254→V | 66, 71 |
| · | 255 | V | V255→I | 14, 178, 203 |
| · | 265 | · | V265→I | 10, 40, 180 |
| · | 375 | L | L375→M | 9, 53, 79, 91, 127 |
| · | 418 | V | V418→A | 14, 24, 139 |
| · | 449 | C | A449→S | 154, 165 |
| · | · | · | C449→A | 53 |
| · | · | · | C449→S | 10, 32, 66, 71, 114 |
| · | · | · | S449→A | 93 |
| · | 471 | V | V471→A | 10, 14, 19, 53, 180 |

**Table S3**—*cont.*
| Genus | Site | Amino acid<br>at the root of<br>phylogeny | Substitution | Branch numbers |
| --- | --- | --- | --- | --- |
| *Viburnum* | 95 | S | S95→T | 107, 144, 25 |
| . | 219 | L | L219→V | 152, 262, 31, 33 |
| . | 225 | I | I225→L | 123, 277, 135, 147, 250, 35 |
| . | 226 | Y | Y226→F | 135, 276, 224, 228 |
| . | 249 | E | E249→D | 75, 81 |
| . | 251 | M | M251→I | 121, 227, 290, 68 |
| . | 255 | V | V255→I | 135, 152, 242, 16, 92 |
| . | 262 | . | V262→A | 135, 276, 92 |
| . | 279 | T | T279→S | 121, 227, 33, 68, 76, 92 |
| . | 309 | M | M309→I | 121, 229, 68 |
| . | 317 | A | A317→G | 135, 152, 276, 14 |
| . | 328 | . | A328→S | 121, 145, 187, 198, 68 |
| . | 340 | E | E340→D | 160, 174 |
| . | 449 | C | A449→S | 0 |
| . | . | . | C449→S | 261, 242, 289 |
| . | 461 | V | V461→I | 14, 92 |
| . | 472 | M | M472→V | 99, 276 |

**Table S 4.** Tests for correlation of amino acid substitution with biome shifts in *Viburnum*. For each site, models were fit to test independent evolution of amino acid and biome, amino acid transitions dependent on biome transitions, and biome transitions dependent on amino acid transitions. The model with the lower Akaike Information Criterion (AIC) out of the latter two models was chosen to be compared to the first model with a likelihood ratio test and shown in this table. ΔAIC, difference in AIC between dependent and independent models; *p*, *p*-value of likelihood ratio test.

| Site | Substitution | ΔAIC | $p$ | Dependent variable | State favored by derived amino acid |
| --- | --- | --- | --- | --- | --- |
| 76 | S76→N | -2.8 | 0.034 | climate | non-cold temperate |
| 95 | S95→T | -3.9 | 0.016 | amino acid | symmetrical |
| 157 | V157→S | -6.2 | 0.006 | amino acid | non-cold temperate |
| 219 | L219→C | -4.3 | 0.012 | climate | symmetrical |
| · | L219→V | -0.8 | 0.095 |  |  |
| 225 | I225→L | -6.5 | 0.004 | climate | symmetrical |
| 226 | Y226→F | -2.2 | 0.039 | climate | symmetrical |
| 230 | A230→T | 3.3 | 0.708 |  |  |
| 249 | E249→D | -0.8 | 0.096 |  |  |
| 251 | M251→I | -2.1 | 0.044 | amino acid | symmetrical |
| 255 | V255→I | -4.5 | 0.014 | climate | non-cold temperate |
| 262 | V262→A | -2.6 | 0.033 | climate | symmetrical |
| · | V262→T | 1.9 | 0.349 |  |  |
| 279 | T279→S | 3.2 | 0.671 |  |  |
| 309 | M309→I | -5.2 | 0.010 | climate | cold temperate |
| 317 | A317→G | -1.7 | 0.054 |  |  |
| 328 | A328→G | -0.2 | 0.136 |  |  |
| · | A328→S | -3.6 | 0.023 | amino acid | non-cold temperate |
| 340 | E340→D | -0.4 | 0.125 |  |  |
| 449 | C449→A | 0.8 | 0.203 |  |  |
| · | C449→S | 1.5 | 0.460 |  |  |
| 461 | V461→I | 2.0 | 0.886 |  |  |
| 472 | M472→V | -2.7 | 0.030 | climate | symmetrical |
| 475 | L475→I | 2.0 | 0.865 |  |  |

**Table S 5.**
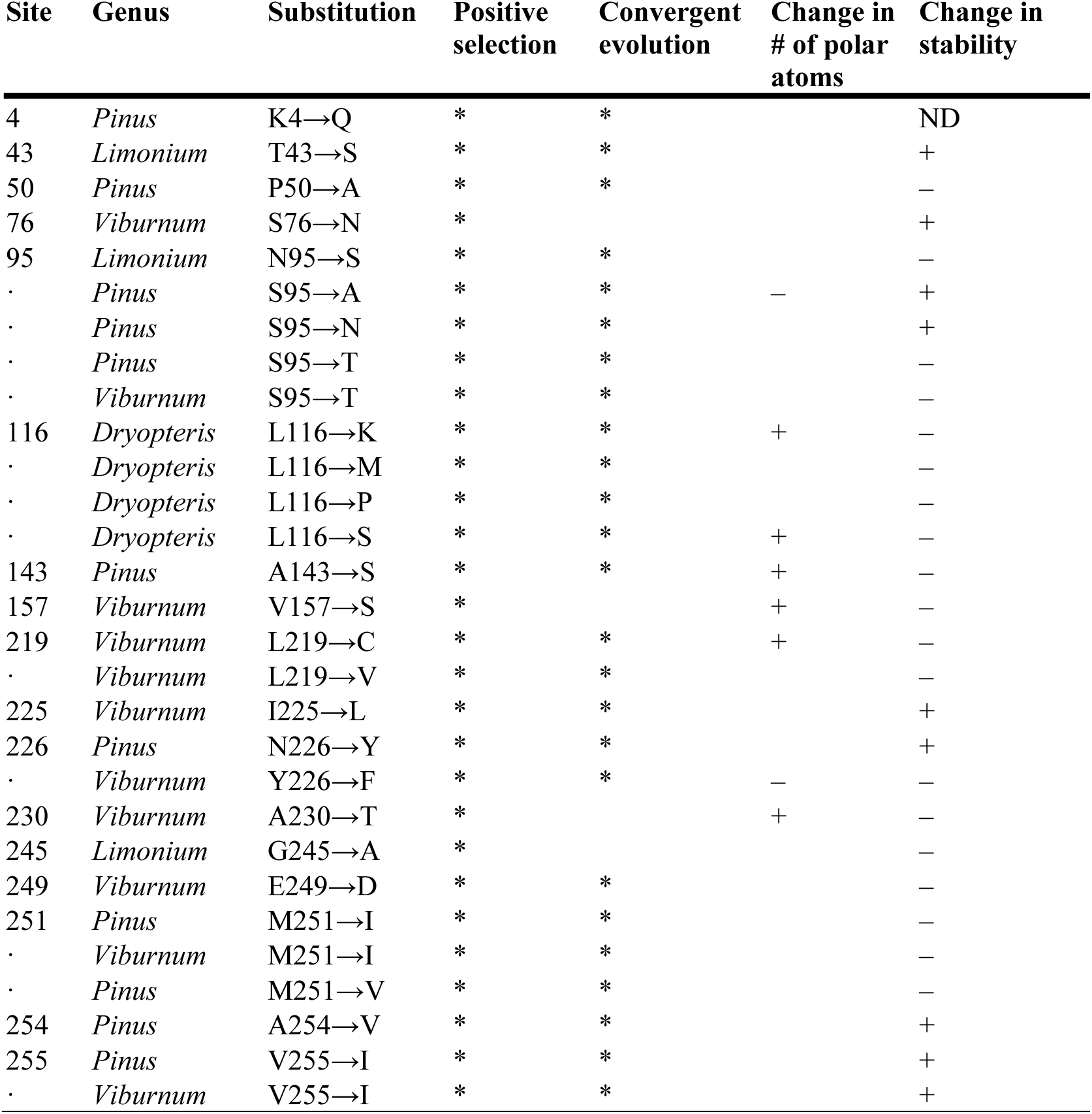
Summary of structure, stability, and phylogenetic analyses for all sites under positive selection or convergent evolution. *, positive selection or convergent evolution with posterior probability > 0.95; +, increase in H-bond ability or increase in stability; –, decrease in H-bond ability or decrease in stability. ND, not determined because the amino acid site is not represented in the crystal structure.

**Figure S 1.**
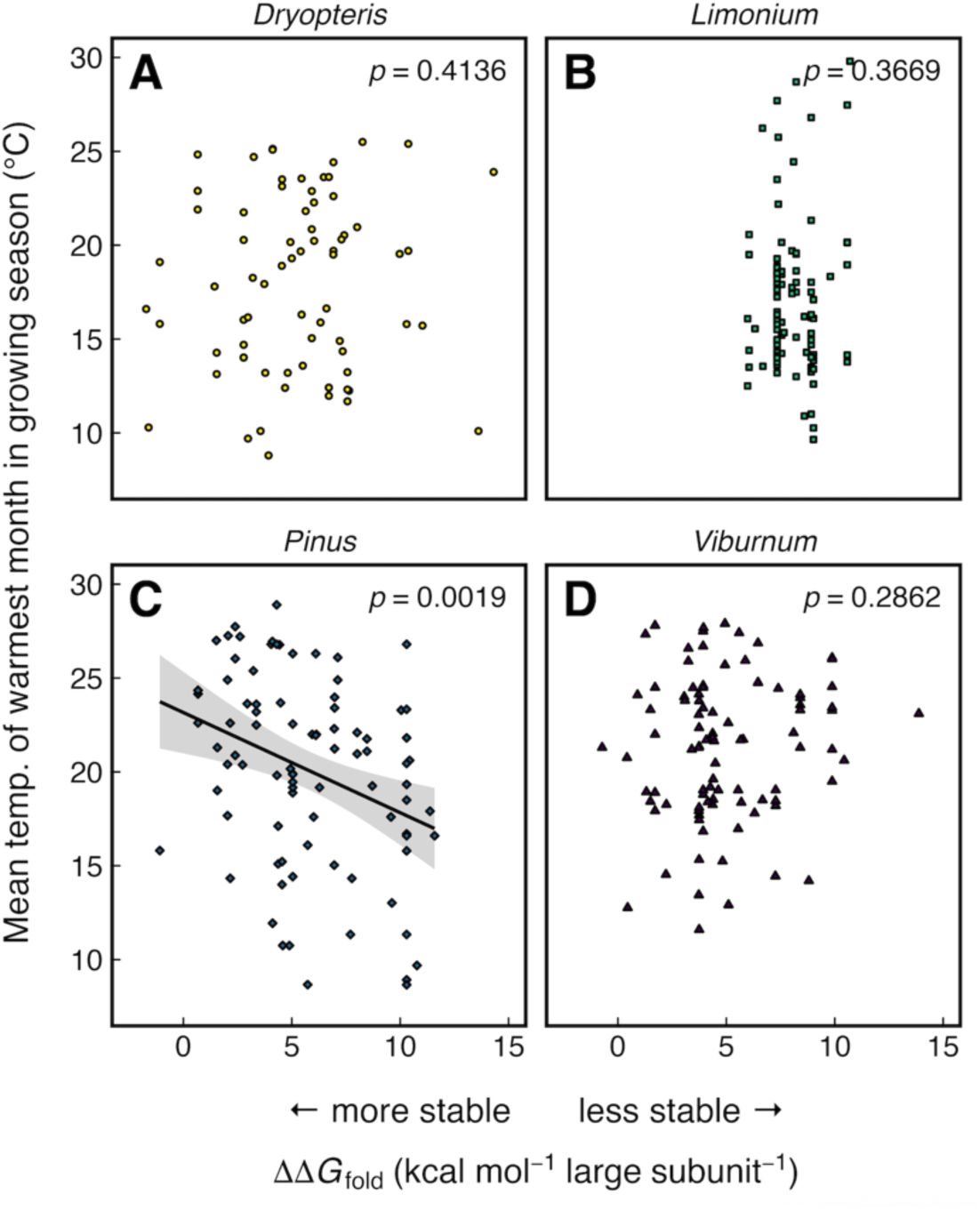
Correlations between growing season temperature and change in free energy of folding (ΔΔ*G*_fold_) for each genus (*n* = 93, 110, 87, and 100, respectively, for *Dryopteris*, *Limonium*, *Pinus*, and *Viburnum*). The solid line is a linear regression, and the grey shaded area represents the 95% confidence interval.

